# An integrated genomics approach towards deciphering human genome codes shaping HIV-1 proviral transcription and fate

**DOI:** 10.1101/2020.05.01.072231

**Authors:** Holly Ruess, Jeon Lee, Carlos Guzman, Venkat Malladi, Iván D’Orso

**Affiliations:** Bioinformatics Core Facility, The University of Texas Southwestern Medical Center, Dallas, TX 75390; Department of Microbiology, The University of Texas Southwestern Medical Center, Dallas, TX 75390; Bioinformatics and Systems Biology Graduate Program, University of California San Diego, La Jolla, CA 92093

**Keywords:** HIV-1, provirus, integration, orientation, transcription, human genome, genomics, machine learning, enhancers, sub-compartments, ChromHMM

## Abstract

A large body of work has revealed fundamental principles of HIV-1 integration into the human genome. However, the effect of the integration site to proviral transcription activity has so far remained elusive. Here we combine open-source, large-scale datasets including epigenetics, transcriptome, and 3D genome architecture to interrogate the chromatin states, transcription activity landscape, and nuclear sub-compartments around HIV-1 integration sites in CD4^+^ T cells to decipher human genome codes shaping the transcription of proviral classes defined based on their position and orientation in the genome. Using a Hidden Markov Model, we describe the importance of specific chromatin states and genome architecture in the control of HIV-1 transcription activity. Additionally, implementation of a machine-learning logistic regression model reveals upstream chromatin accessibility, transcription activity, and categorical nuclear sub-compartments as optimal features predicting HIV-1 transcriptional outcomes. We finally demonstrate clinical relevance by interrogating the positions of intact proviruses persisting in patients under suppressive therapy and provide a compass compatible with clinical decision-making.

## INTRODUCTION

One of the most exciting breakthroughs in biomedical research was the discovery of highly suppressive anti-retroviral therapy (ART), which curbs replication of human immunodeficiency virus type 1 (here referred to as HIV) to nearly undetectable levels (Perelson et al. 1997). However, ART does not eliminate the transcriptionally silent, yet replication-competent latent provirus, which persist indefinitely despite highly suppressive therapy (Chun et al. 1995; Chun et al. 1997; Chun et al. 1998) and can be reactivated upon therapy cessation leading to rebound of plasma viremia (Chun et al. 2010; Wen et al. 2018).

Over the past two decades, great efforts have been made for elucidating how HIV integrates into the human genome and how HIV proviral transcription normally operates (Ott et al. 2011; Mbonye and Karn 2014; Hughes and Coffin 2016; Mbonye and Karn 2017; Michieletto et al. 2019; Morton et al. 2019). Interestingly, in CD4^+^ T cells of patient samples and in *ex vivo* studies, HIV preferentially integrates into chromatin-accessible sites located near the nuclear pore and within or near transcriptionally active regions (Schroder et al. 2002; Cohn et al. 2015; Marini et al. 2015; Lucic et al. 2019). However, integrated proviruses are detected on every human chromosome, in various chromatin landscapes (euchromatic and heterochromatic), and at different locations (intergenic or intragenic) and orientations (sense, divergent or convergent) respective to human genes and regulatory elements. Since the integration landscape is highly diverse in sequence and chromatin structure (Bukrinsky et al. 1991; Jordan et al. 2001; Sherrill-Mix et al. 2013; Battivelli et al. 2018), it seems logical to speculate that the integration site neighborhood could contain information influencing the magnitude of transcription thereby shaping proviral fate (future viral replication: active vs latent). Previous work on several Jurkat CD4^+^ T cell clones containing individual HIV integrants revealed that the integration site controls basal and immune stimulation-dependent transcription (Jordan et al. 2001), indicating that HIV possibly operates in an integration site-dependent manner and not cell-autonomously. Despite these past discoveries, it remains unknown what genomic features or combination of them control proviral transcription and fate.

It has been known for years that the human genome provides the underlying code for the correct transcriptional regulation of biological processes through precise spatial-temporal interactions between *cis*-elements present at promoters and enhancers, and sequence-specific factors that recognize them, as well as by the position of genes and regulatory elements in the three-dimensional (3D) space and in relation to nuclear territories (Heintzman et al. 2007; Dixon et al. 2012; Rao et al. 2014; Dekker and Mirny 2016; Andersson and Sandelin 2020). These regulatory elements could provide local or distal functions thereby influencing HIV proviral transcription and expression. Given these ideas, we hypothesized that the integration site contains “instructions” – here referred to as integration code (one or more genomic features located proximally and/or distally from the integration site) – that shape the organization and activity of proviruses in the context of the human genome.

Here we use an integrated genomics approach, interrogate patient samples, and implement a machine learning logistic regression model to both define and predict molecular features contributing to proviral expression and persistence (Fig. 1). First, by combining open source, large-scale datasets including epigenetics, transcriptome, and 3D genome architecture (Supplemental Table S1), we defined the chromatin states, transcription activity landscape, and nuclear sub-compartments around HIV integration sites in CD4^+^ T cells and provide evidence of the importance of HIV placement on the human genome to its transcriptional activity. Second, by implementing a machine learning logistic regression model of a 2-kb region around HIV integration sites, we predict upstream chromatin accessibility, transcription activity, and categorical nuclear sub-compartments as optimal features shaping HIV transcriptional outcomes. Third, by interrogating patient samples bearing intact proviruses persisting under suppressive therapy we demonstrate clinical relevance and propose guidelines for future interventions.

**Figure 1.**
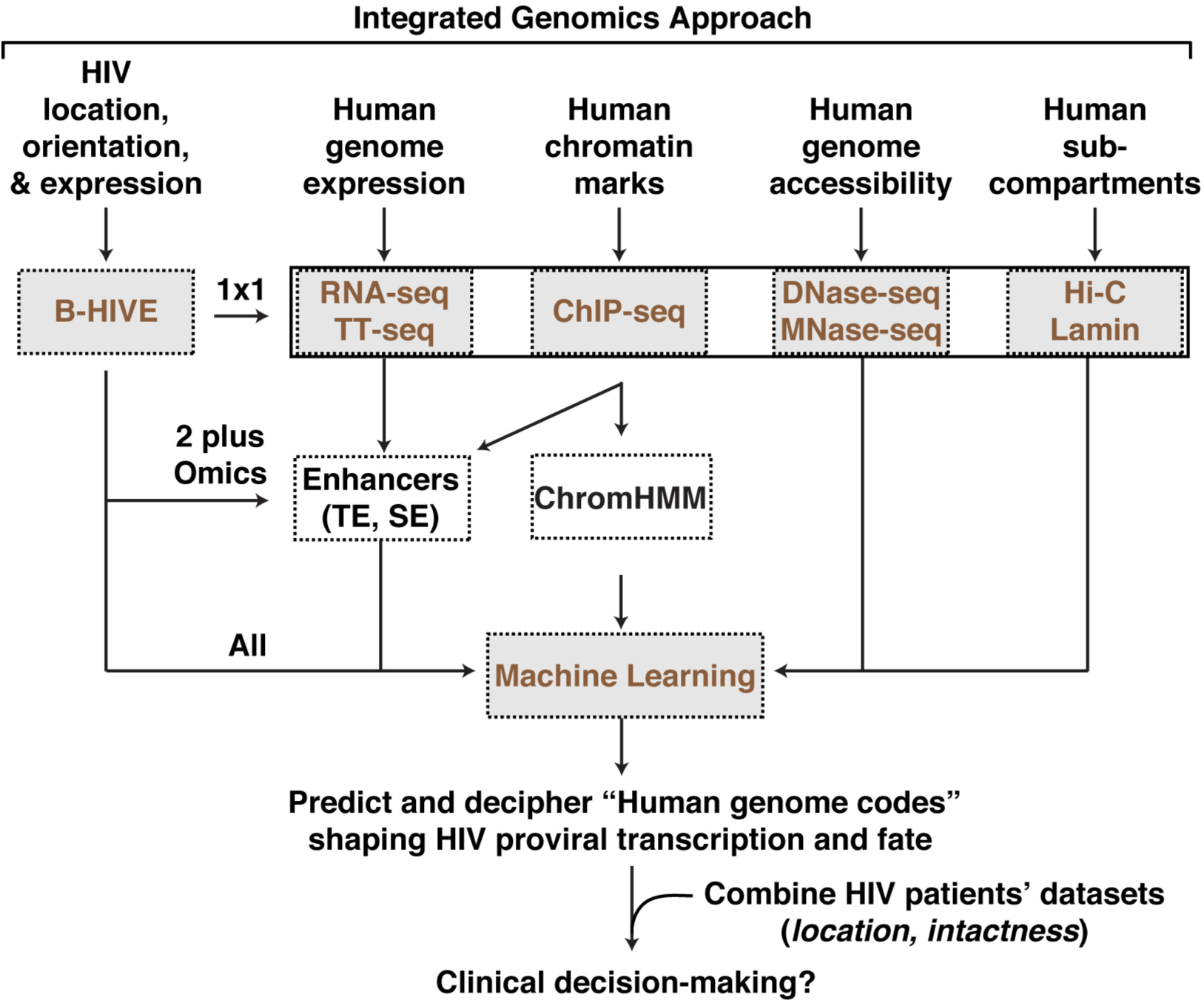
Flow chart of the integrated genomics approach to elucidate human genome codes contributing to HIV proviral transcription and fate. B-HIVE expression data is compared to each individual dataset one-by-one (“1×1”). Then, B-HIVE expression data is compared to datasets that combine multiple datasets (“2 plus Omics”) (e.g., typical enhancers (TE) and super enhancers (SE) combine TT-seq and ChIP-seq). B-HIVE is then compared to “all” datasets with a Machine Learning model. From there, we then integrated HIV patients’ datasets with the long-term objective to predict human genome codes that can lead to clinical decision-making.

## RESULTS

### Expression of HIV Integration Groups Defined Based on Their Position and Orientation Respective to Genes in the Human Genome

To start interrogating the relationship between HIV integration position and transcription activity, we first reanalyzed the B-HIVE dataset in Jurkat CD4^+^ T cells (Chen et al. 2017), which used the thousands of reporters integrated in parallel (TRIP) assay (Akhtar et al. 2013), to obtain HIV position information (integration site) respective to human genome coordinates. Because HIV integrates within (intragenic) or between (intergenic) genes, and in the same or opposite orientation respective to the nearest human gene transcription start site (TSS) (Fig. 2A), we first defined 6 integration groups based on their positions and orientations to explore any potential relationships of each group with human genomic features. Of note, Group 6 is a composite of 3 subgroups depending on the three possible combinations of HIV directions respective to the two overlapping genes [Group 6a: both genes in same direction with HIV in same direction (n=26); Group 6b: gene 1 and gene 2 in opposite directions (n=24); and Group 6c: both genes in same direction with HIV in opposite direction (n=10)] (Fig. 2A). However, given the limited number of genes in each of the subgroups of Group 6, we treated them as a single group to increase statistical power. Remarkably, the large number of proviruses detected in Groups 4-6 (Intragenic) compared to Groups 1-3 (Intergenic) is consistent with previous studies examining integration sites *ex vivo* and in patient samples, highlighting HIV integrations in intronic regions of highly transcribed genes (Schroder et al. 2002; Ikeda et al. 2007; Maldarelli et al. 2014; Lucic et al. 2019).

**Figure 2.**
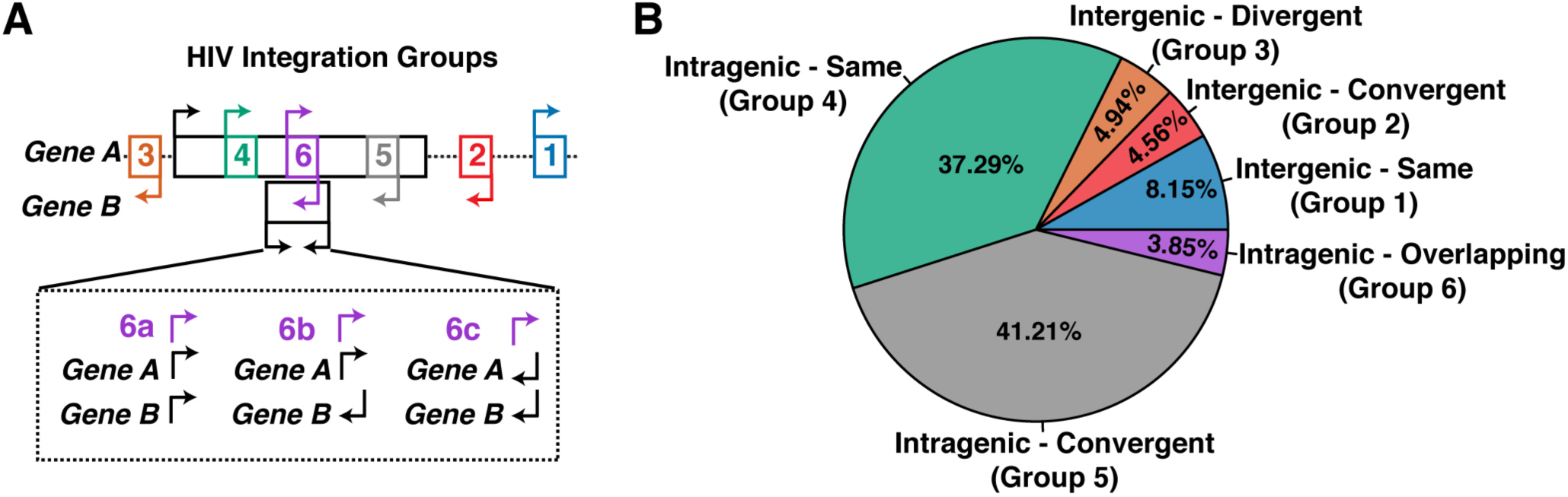
Defining the expression of HIV proviral integration groups based on their position and orientation respective to human genes. (A) Diagram of 6 HIV integration groups relative to nearest gene(s). Group 6 (overlapping genes at the same position) is comprised of three subgroups (6a, 6b, 6c). (B) Pie chart of the percentage of B-HIVE insertions (n = 1558) into each of the 6 HIV integration groups.

Having established the HIV integration groups, we then examined the relationship between HIV positions and their expression (see Methods). After visualizing the expression of each HIV integration site on a per integration group basis using Circos plots, we found that HIV proviruses from each group are detected in every single chromosome with various expression levels irrespective of their group (Supplemental Fig. S1A-F). Together, this indicates that the arrangement of insertions in the position/orientation-defined groups above is not the only determinant of HIV expression levels. Therefore, we reasoned that HIV proviral transcription activity might be regulated by local and/or distal codes that are unique to the integration site and not shared within each integration group.

### Relationship Between HIV Expression and Distance to Nearest Human Gene TSS

Given that proviruses can be located at various distances respective to nearby gene TSS, we interrogated if there is any relationship between the expression level of proviruses in each HIV integration group and their distance to nearby TSS to test the hypothesis that proviruses located closer to TSS are more active than those located at farther distances (Fig. 3A).

**Figure 3.**
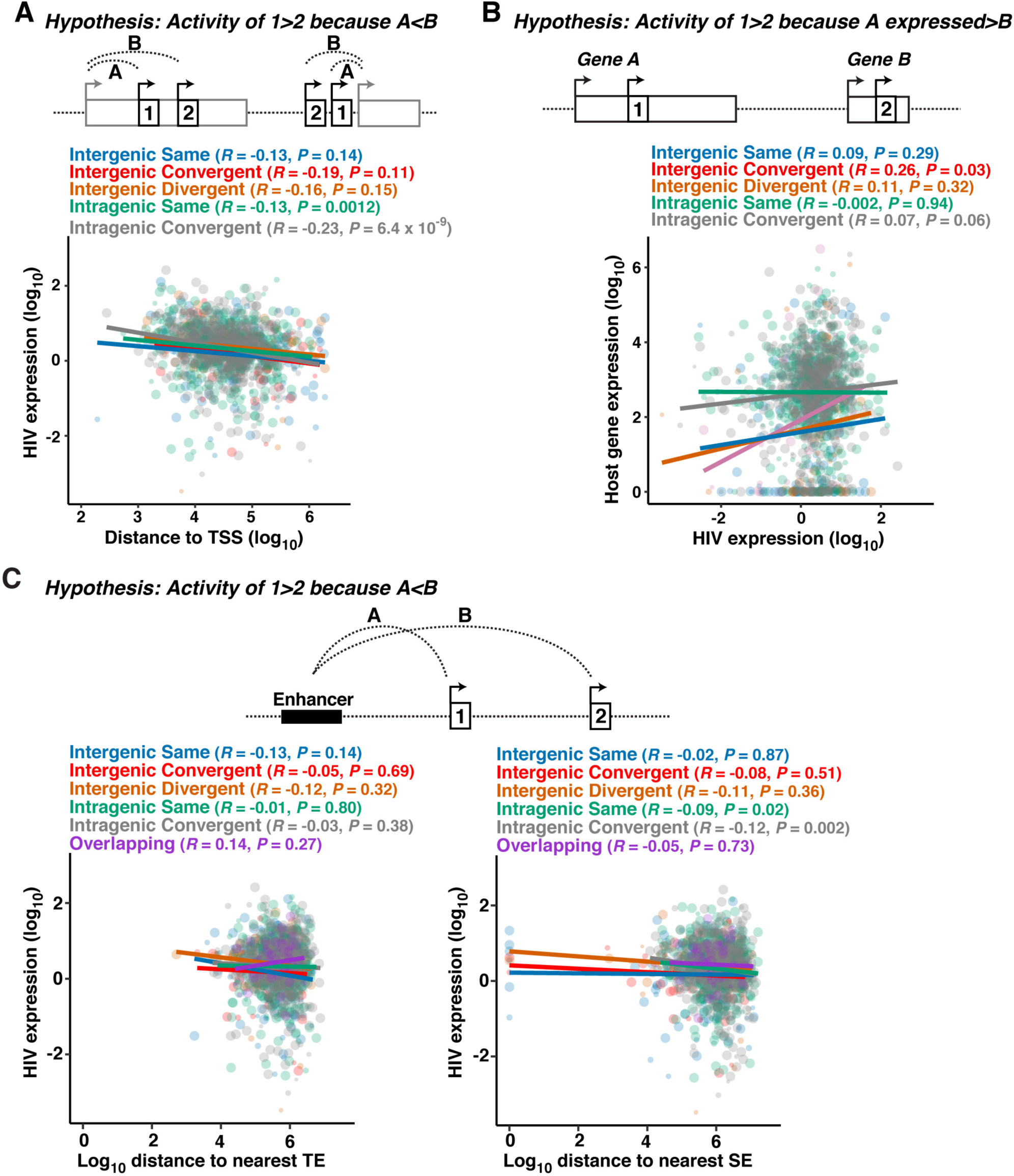
Scatter plots of the Pearson correlation coefficient (*R*) and p-value (*P*) between proviral expression of each HIV integration group to nearest gene TSS, gene expression, and enhancers. (A) Hypothesis that HIV expression is stronger the closer it is to the nearest gene TSS. Correlation of HIV expression (log_10_) as a function of distance to TSS (log_10_) for each of the 5 HIV integration groups. (B) Hypothesis that HIV expression is correlated to the expression of its nearest gene, where if the nearest gene has higher expression, HIV would also have higher expression. The opposite is hypothesized for lower expressed genes. Correlation of HIV expression (log_10_) as a function of host gene expression (log_10_) for each of the 5 HIV integration groups. (C) Hypothesis that HIV expression is higher the closer it is to an enhancer. Correlation of HIV expression (log_10_) to the log_10_ distance of the nearest typical enhancer (TE) or super enhancer (SE).

Interestingly, we found that the expression of proviruses in intergenic regions is poorly correlated with the distance to the nearest TSS irrespective of their orientation [Group 1: Intergenic - Same (*R* = - 0.13, *P* = 0.14), Group 2: Intergenic - Convergent (*R* = −0.19, *P* = 0.11), and Group 3: Intergenic - Divergent (*R* = −0.16, *P* = 0.15)] (Fig. 3A), potentially indicating that the orientation of intergenic proviruses is not a major feature controlling HIV expression. For this analysis we did not include Group 6 because these HIV insertions are within two overlapping genes, which either of the genes or both genes could influence HIV expression, leading to uncertainty of the analysis.

However, we observed a potential pattern for intragenic proviruses in Groups 4-5 (Group 4: Intragenic – Same (*R* = −0.13, *P* = 0.0012); Group 5: Intragenic – Convergent (*R* = −0.23, *P* = 6.4×10^−9^)), which seem to be, in general, more significantly correlated than the intergenic proviruses in Groups 1-3 (Fig. 3A). Surprisingly, the expression of proviruses in Group 5 was more correlated with the distance to TSS of its nearby gene than the expression of proviruses in Group 4, suggesting that the orientation of intragenic HIV proviruses respective to the nearest TSS impacts proviral expression, in agreement with theories of transcription interference of proviruses positioned in the same orientation as its associated genes (Lenasi et al. 2008).

### Relationship Between HIV Activity and the Expression of its Associated Human Gene

Another regulatory feature shaping HIV proviral transcription can be the level of expression of the HIV-associated gene. Thus, it is possible that the activity of provirus 1 is greater than provirus 2 if the gene associated with provirus 1 (Gene A) is expressed at higher levels than the gene linked to provirus 2 (Gene B) (Fig. 3B). To test this hypothesis, we calculated the expression of proviral Groups 1-5 (Fig. 2A) and the expression of the HIV-associated human gene. Once again, we excluded Group 6 (Overlapping) because of the complexity of HIV association with two overlapping genes. Surprisingly, we found no apparent correlations for Group 1: Intergenic - Same (*R* = 0.09, *P* = 0.29) and Group 3: Intergenic - Divergent (*R* = 0.11, *P* = 0.32). However, Group 2: Intergenic - Convergent, showed a slight, but statistically significant correlation (*R* = 0.26, *P* = 0.03), potentially indicating that this HIV arrangement somehow benefits from host transcription activity.

Given HIV preferably integrates inside genes with no apparent difference in orientation respective to host gene transcription activity, we were particularly interested in testing whether there is any correlation between the expression of HIV in Group 4: Intragenic - Same, and Group 5: Intragenic - Convergent. Strikingly, we found that the correlation level of Group 4: Intragenic - Same (*R* = −0.002, *P* = 0.94) is much lower than the correlation of Group 5: Intragenic - Convergent (*R* = 0.07, *P* = 0.06), which is trending towards statistical significance (Fig. 3B). These results suggest that the convergent orientation offers HIV a molecular benefit potentially linked to the lack of transcription interference by RNA polymerase II molecules transcribing host genes positioned in the same orientation as HIV (Lenasi et al. 2008), consistent with the better correlation of expression of this group and the distance to nearest TSS (Fig. 3A).

Taken together, our large-scale data analysis indicates that transcription interference accounts for at least part of the observed proviral expression effects. However, since the correlations are low, it is likely that a combination between orientation respective to nearest gene promoters and other regulatory features account for HIV expression levels. Thus, below we also interrogate HIV expression as a function of its position in relation to enhancers, nuclear sub-compartments, and chromatin states to provide an integrated analysis (as described in Fig. 1).

### Contribution of Human Genome Enhancers to HIV Proviral Transcription

HIV displays specific preference to integrate into genes proximal to high density of enhancers (Lucic et al. 2019), which can potentially regulate proviral transcription. Enhancers are short DNA sequences that act as transcription factor binding hubs controlling key transcriptional programs by fine-tuning target gene promoter activity across vast linear distances (Whyte et al. 2013; Kim and Shiekhattar 2015; Li et al. 2016). Given their importance, we explored whether the position of HIV respective to enhancers is a key regulatory element for determining proviral activity level. To this end, we generated a rigorous and comprehensive database of active enhancers based on a combination of conventional epigenetic and transcription signatures including: 1) a unique chromatin state demarcated by high H3K27ac, high H3K4me1, and low H3K4me3 derived from ChIP-seq, and 2) symmetrical bi-directional enhancer RNA transcription (eRNA) derived from TT-seq (Supplemental Fig. S2A). At least two types of active enhancer classes have been described: typical enhancers (TE) and super enhancers (SE). TE contain the classic composition of features indicated above, whereas SE are locally grouped clusters of enhancers (defined as a higher signal of H3K27ac, H3K4me3, H3K4me1, and active transcription (TT-seq)) domains within 12.5-kb of each other (Supplemental Fig. S2A) driving high levels of transcription of nearby cell-identity genes (Whyte et al. 2013).

A Hidden Markov Model (HMM) of eRNA Watson and Crick strands was used to classify transcribed regions that when overlapped with known enhancer histone marks identified 2,180 active intergenic TE (Supplemental Fig. S2B-C). We defrayed from identifying intragenic enhancers given the high content of genic transcription and histone marks potentially obscuring accurate identification of this class of enhancers. Using this approach, we also identified 767 SE containing the conventional high density of clustered H3K27ac, H3K4me3, H3K4me1, and TT-seq activity (Supplemental Fig. S2A-C).

To test the hypothesis that HIV proviral expression is correlated to its proximity to enhancers (Fig. 3C) we used the rigorously assembled enhancer database to measure both the expression and distance of each provirus of the 6 different HIV integration groups to the nearest TE and observed poor correlation and no statistical significance (*P* < 0.05) for all groups, suggesting that proximity to a TE alone is not a good predictor of HIV proviral activity (Fig. 3C).

Given previous reports that genes located near SE are transcribed to much higher levels compared to genes located near TE (Whyte et al. 2013), we hypothesized that if the proximity to enhancers is a true regulator of proviral transcription we would then expect that, compared to less active or latent proviruses, the most active proviruses should be positioned nearer to SE. Therefore, we also evaluated the distance of each provirus of the 6 HIV integration groups to the nearest SE and found low correlations and no statistical significance for all three intergenic groups (Groups 1-3: same, convergent, divergent, respectively) (Fig. 3C). However, interestingly, the intragenic groups (Groups 4-5) showed a statistical significant correlation (Group 4: Intragenic - Same (*R* = −0.09, *P* = 0.02) and Group 5: Intragenic - Convergent (*R* = −0.12, *P* = 0.002)) (Fig. 3C), consistent with the better correlations between the expression of both intragenic groups and the distance to the nearest gene TSS (Fig. 3A) and transcription activity (Fig. 3B), thus proposing that intragenically positioned proviruses may benefit from both its positions nearby to regulatory elements such as gene promoters and SE. Together, the preference for HIV to integrate into genes proximal to SE (Lucic et al. 2019) may have the dual benefit of increasing proviral transcriptional levels.

### Contribution of Chromatin States to HIV Proviral Transcription

Since chromatin marks do not denote a singular function, and because we wanted to classify regions of the genome to precisely define chromatin states to explore their contribution to HIV proviral transcription (Fig. 1), we implemented a 15-state HMM derived from ChromHMM (Ernst and Kellis 2017) using 7 histone marks to annotate chromatin states similar to the annotations from the Roadmap Epigenomics Project (Roadmap Epigenomics et al. 2015). To *de novo* generate the core 15-state model in Jurkat T cells we compared the relative abundance of the state emissions in Jurkat T cells with known chromatin states for the 3 ENCODE cell lines most genetically and phenotypically linked with Jurkat (E115: Dnd41 T cell leukemia, E116: GM1282878 lymphoblastoid, and E123: K562 T cell leukemia). After overlapping the Jurkat T cell states with these 3 related epigenomes and comparing each histone mark function, we rearranged and relabeled the state numbers to correspond to identical labels from Roadmap Epigenomics Project (Fig. 4A; Supplemental Fig. S3A-C).

**Figure 4.**
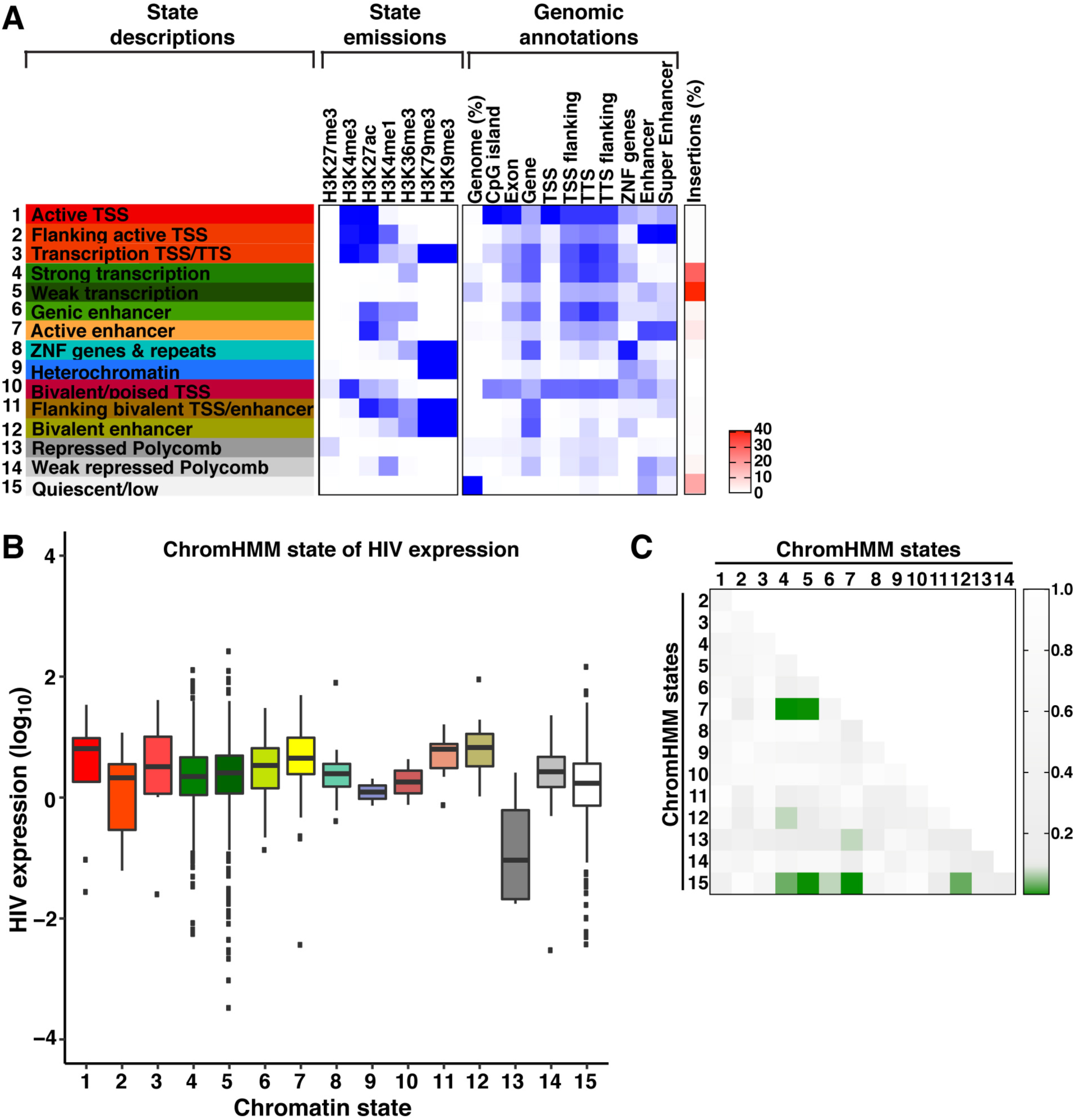
Expression of HIV proviruses in relation to chromatin states defined using ChromHMM. (A) From left to right: ChromHMM plot containing the 15 states. Histone marks used to create the states (state emissions). Overlap enrichment plots (ChromHMM) of RefSeq genomic annotations, TE (Jurkat), and SE (Jurkat) by 15 states. Heatmap of percentage of insertions in each state. (B) Box plot of B-HIVE expression by 15-state model. (C) Heatmap of *P* values comparing the median expression of B-HIVE in 2 different states by Kruskal-Wallis rank sum test.

Given that HIV integrates within and/or near open chromatin based on accessibility and epigenetic data (Lucic et al. 2019), we would expect HIV to be inserted into open chromatin states. As expected, HIV was more likely to insert into “accessible” chromatin (states 2-8, 11-12, and 14 (*P* < 0.05, two-proportions z-test)) and less likely to insert into “inaccessible” chromatin (states 13 and 15 (*P* < 0.001, two-proportions z-test)) (Supplemental Fig. S4).

Since HIV is more likely to insert into accessible regions of the genome, we also wanted to compare proviral expression in each chromatin state to determine if the median expression between states is different. HIV expression was not found to be normally distributed or have similar variances between states, so a Kruskal-Wallis rank sum test computed significant differences of the median between different states. Surprisingly, HIV expression was higher in state 7 (Active enhancer) than states 4 or 5 (Strong and Weak transcription, respectively, *P* < 0.05) (Fig. 4B,C). Not surprisingly, state 15 (Quiescent/Low) has lower expression than states 4, 5, 7, and 12 (Strong transcription, Weak transcription, Active enhancer, and Bivalent enhancer, respectively, *P* < 0.05) (Fig. 4B,C).

Together these results reveal that HIV is more likely to integrate into accessible regions of the genome (Supplemental Fig. S4), and that open regions associated with promoter and enhancer activity have higher HIV mean expressions than those in inaccessible regions.

### Contribution of Nuclear Spatial Sub-compartments to HIV Proviral Transcription

The 3D organization of chromosomes enables long-range interactions between enhancers and promoters that are critical for building complex gene regulatory networks (Dekker and Mirny 2016; Yu and Ren 2017). Interphase chromosomes occupy separate spaces known as nuclear territories (Meaburn and Misteli 2007), and each chromosome is organized into dynamic, non-random structures containing stretches of transcriptionally active compartments interspersed with sections of transcriptionally inactive compartments (Lieberman-Aiden et al. 2009). As such, the genome is partitioned into contact domains (A and B) segregating into 6 sub-compartments (A1, A2, B1, B2, B3, and B4) that: 1) appear located in different nuclear territories, 2) are associated with distinct patterns of histone marks, and 3) show different expression levels (Rao et al. 2014). To examine both the distribution and expression of HIV proviruses in the various sub-compartments we utilized two complementary approaches. First, we re-analyzed a Hi-C dataset in Jurkat T cells (Lucic et al. 2019), and second, we predicted sub-compartments in Jurkat T cells using the lamin sub-compartments derived from GM12878 cells (Rao et al. 2014).

Using the Jurkat Hi-C dataset, we identified the A and B sub-compartments using the first eigenvector. We then further divided the A compartment into A1 and A2 using a k-means clustering of k = 2. However, k-means clustering of the B compartment, with k = 2 through k = 5, did not yield a clearly defined separation of three regions based on expected chromatin marks (Supplemental Fig. S5), leaving us to keep all B sub-compartments together. The distribution of proviruses revealed there is a preferential insertion into the A sub-compartments (*P* < 0.001, two-proportions z-test), and reduced insertions within B sub-compartment (*P* < 0.001, two-proportions z-test), than by random occurrence (Supplemental Fig. S6A,C) given the Jurkat T cell sub-compartments coverage (Supplemental Fig. S6B), consistent with recent observations (Lucic et al. 2019). The mean HIV expressions in the A1 and A2 sub-compartments was significantly higher from the B compartment (*P* < 0.001, Kruskal-Wallis rank sum test) without any obvious, statistical significant difference between the A1 and A2 sub-compartments (*P* = 0.43) (Supplemental Fig. S6D), suggesting that HIV integration into more accessible regions of the genome is, in general, beneficial for proviral expression.

Since we noticed the Jurkat Hi-C dataset might be too sparse to divide into 6 sub-compartments, we then used the higher resolution sub-compartments defined in GM12878 cells. GM12878 is a T cell line genotypically and phenotypically closely related to Jurkat (Rao et al. 2014), thus arguing that these results could be applicable to Jurkat, since domains are typically conserved between cell types. Using this data, we found a significant increase of HIV proviruses in sub-compartments A1, A2, and B4, and significantly lower insertions in sub-compartments B1, B2, B3 (*P* < 0.001, two-proportions z-test) (Supplemental Fig. S6E-G). Also, there is an enrichment of mean HIV expression in sub-compartments A1 vs A2, B1, B2 or B3 (*P* < 0.001, Kruskal-Wallis rank sum test) (Supplemental Fig. S6G,H), suggesting the location of HIV into categorical nuclear sub-compartments overall contributes to increased expression. Notably, these results are similar to the Hi-C data analysis in Jurkat cells, suggesting there is a conservation of sub-compartments between the two cell types, consistent with previous results (Rao et al. 2014).

### Machine Learning Approach to Train a Model Predicting HIV Proviral Transcription and Fate

To foresee features and guide experimental approaches to clinical analysis, a machine learning approach was used to train a logistic regression model for HIV expression prediction. We examined 2-kb regions, in 200-bp increments, around HIV integration sites (Fig. 5A). When a threshold of Information Value (IV) ≥ 2 was applied, 26 features were determined as optimal features for the prediction task, which include all 20 lamin sub-compartment bins, 5 bins in H3K27ac, and 1 bin in MNase-seq (Supplemental Table S2; Supplemental Fig. S7). For the test dataset, HIV expression levels were predicted through the logistic regression model of the optimal features, their estimated weights and corresponding odds ratios, and the standard errors of the estimated weights were obtained (Supplemental Table S3). Notably, the evaluation metrics were calculated as 68.42% of sensitivity, 59.10% of specificity and 64.71% of AUROC, confirming that the model has fair prediction power (Fig. 5B). The predicted HIV expression values for the ‘High’ expression category were found significantly higher than those for the ‘Low’ category (*P* = 0.00025, Wilcoxon test) (Fig. 5C). Meanwhile, the predicted values for the ‘Intermediate’ and ‘Low’ categories showed no significant difference (*P* = 0.27, Wilcoxon test) (Fig. 5C). Furthermore, the actual and predicted HIV expression values were found to have a positive, moderate correlation (*R* = 0.19, *P* = 0.00018) (Fig. 5D).

**Figure 5.**
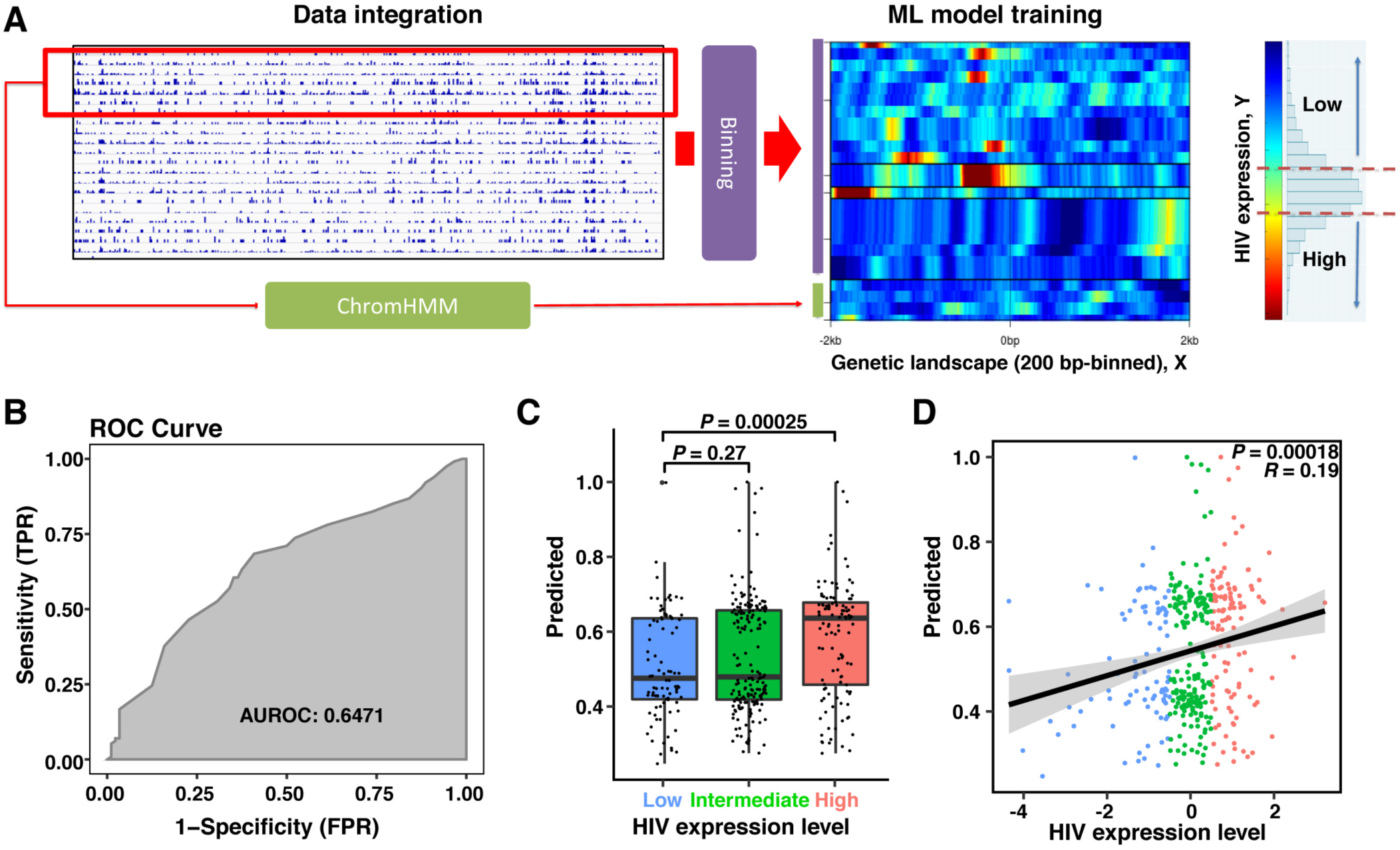
HIV expression level prediction based on genetic and 3D landscape surrounding integration sites. (A) Schematic of data integration and training of the machine learning (ML) model. Genetic marks 2-kb surrounding HIV insertions were binned every 200-bp and measured. B-HIVE data was split into 3 groups (Low, Intermediate, and High), training the ML model for Low versus High. (B) The trained model’s AUROC (Area Under the Receiver Operating Characteristics) curve is shown. (C) Predicted HIV expression comparison among three categories of Low, Intermediate, and High. P-value (*P*) calculated from Wilcoxon test. (D) Linear regression and Pearson correlation test between the actual and predicted HIV expression values.

### Combination of *In Vitro* HIV Integration/Expression Data with Primary and Patient Datasets

To extend our studies to more physiologically relevant models and provide clinical relevance, we first assembled open source, publicly available HIV integration datasets from 11 patients studies (Mack et al. 2003; Han et al. 2004; Ikeda et al. 2007; Maldarelli et al. 2014; Wagner et al. 2014; Kok et al. 2016; Sharaf et al. 2018; Coffin et al. 2019; Einkauf et al. 2019; Garcia-Broncano et al. 2019; McManus et al. 2019) and 4 studies with primary CD4^+^ T cells infected *ex vivo* (Sherrill-Mix et al. 2013; Sunshine et al. 2016; Ferris et al. 2019; Lucic et al. 2019) (Supplemental Table S4). We used these datasets to interrogate the epigenomic landscape of intact vs defective proviruses to provide fundamental principles of differential positioning and regulation of these two proviral classes and to compare their epigenomic landscapes to proviruses in the immortalized (Jurkat T cell) models of latency (Fig. 6A).

**Figure 6.**
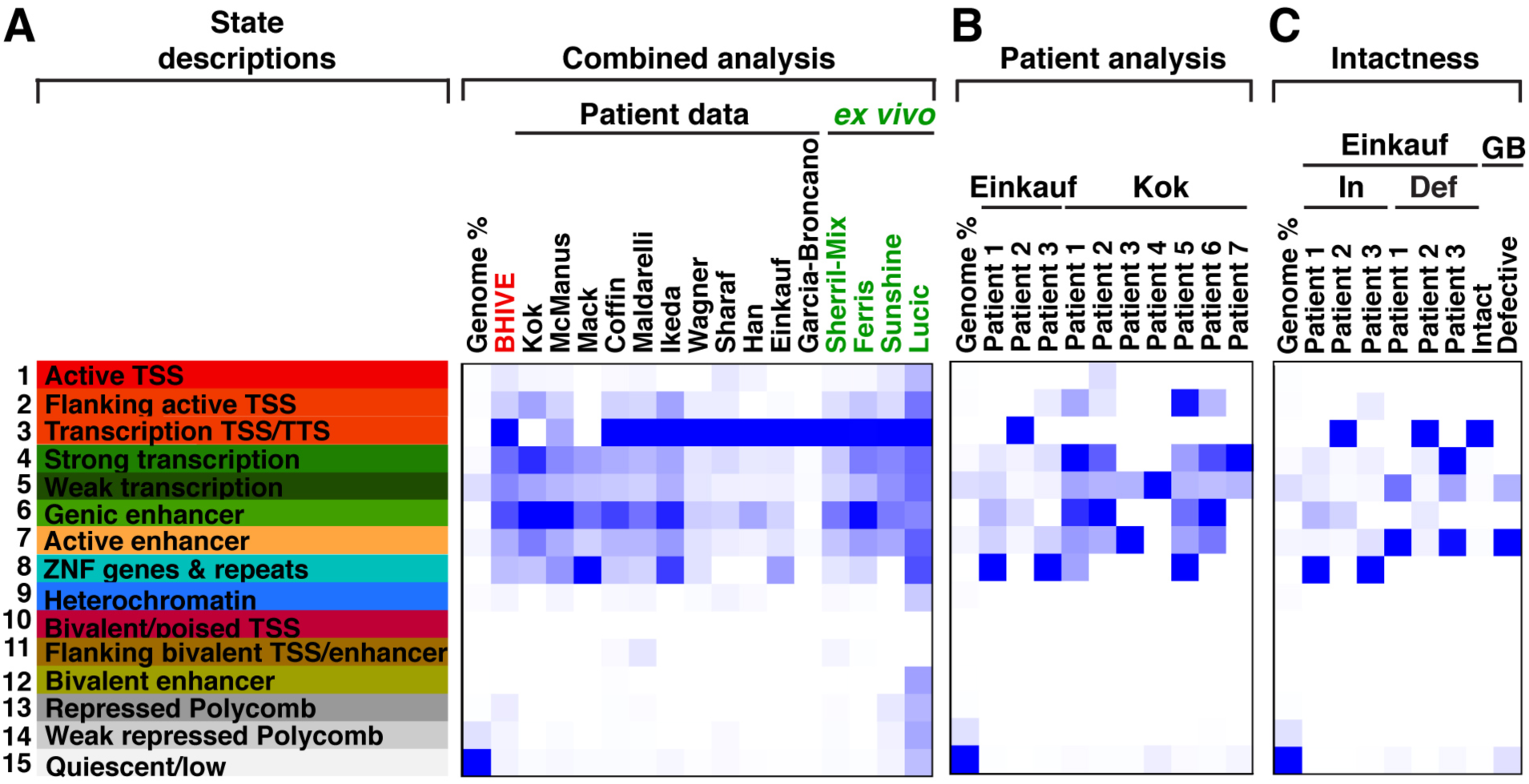
Enrichment of HIV integrants in patient data and their relatedness to B-HIVE in each of the 15 chromatin states. (A) Overlap enrichment plot (ChromHMM) of patient and *ex vivo* data by the 15-state ChromHMM of primary CD4^+^ T helper memory cells (E040). (B) Same as A, but splitting Einkauf and Kok datasets by individual patients. (C) Same as A, but splitting Einkauf patients and Garcia-Broncano (GB) by intact vs defective proviruses.

To accomplish this, we downloaded the chromatin states in 4 primary CD4^+^ T cell states relevant to HIV infection from the Roadmap Epigenomics project: E039 (naïve CD4^+^ T cells), E040 (memory CD4^+^ T cells), E041 (naïve CD4^+^ T cells stimulated with phorbol ester), and E045 (memory and effector CD4^+^ T cells) (Supplementary Fig. S8-S10). From these 4 CD4^+^ T cell states, we choose E040 because it is more relevant to HIV persistence as the virus typically infects activated cells that, over time, persist in resting memory CD4^+^ T cells in patients under suppressive therapy (Shan et al. 2017).

From the combined analysis of patient and *ex vivo* studies, we clearly observed large differences in the integration landscape among datasets and heterogeneous distribution within datasets, with HIV being integrated in the chromatin accessible states (states 1-to-8) but preferentially within states 3, 6, and 8 (Transcription TSS/TTS, Genic enhancer, and ZNF genes & repeats, respectively) (Fig. 6A). Since the percentage of the genome for each chromatin state is not equal, we refer to “preferential” as chromatin states having more HIV integration sites than expected based on the percentage of the genome in each chromatin state. Interestingly, all *ex vivo* datasets and patient datasets except Kok and Mack revealed a preferential integration in state 3 (Transcription TSS/TTS), which, importantly, is also found in the immortalized (B-HIVE) system, demonstrating conserved principles of retroviral integration between immortalized, *ex vivo* and patient studies.

While a few patient datasets showed enrichment of integration positions in state 3 (Transcription TSS/TTS), and much lower levels in state 6 (Genic enhancer) and state 8 (ZNF genes & repeats) such as Einkauf, Wagner, and Han, the majority showed remarkable enrichment in state 4 (Strong transcription), state 5 (Weak transcription) and state 7 (Active enhancer) (Fig. 6A). This result again highlights the heterogeneity in enrichment of the integration landscape, which is most likely a property of retroviral integration that relies on the activation status of the infected cell (Shan et al. 2017). Additionally, a surprising deviation from the commonly seen integration landscape in chromatin accessible states (states 1-8) in all three type of integration datasets (B-HIVE, *ex vivo*, and patient) was seen in the Lucic dataset, which also showed enrichment of integrations in chromatin inaccessible states 12-15 (Bivalent enhancer, Repressed Polycomb, Weak repressed Polycomb, and Quiescent/low, respectively).

Together, not only does this analysis reveal the heterogeneous nature of the HIV proviral integration landscape in various chromatin states and variability among patient datasets, but also exposes commonalities in the integration enrichment patterns between immortalized, primary, and patient datasets. Given this integration pattern conservation and the importance of accessible chromatin states (promoters and SE) in HIV proviral expression in the immortalized (B-HIVE) system, it is likely that the rules defined in the B-HIVE model are conserved in patient samples.

### Analysis of Individual Patient HIV Integrants within Chromatin States

Because the previous analysis was done on the combined integration positions of the various datasets, we then focused on studies bearing information on individual patients. Strikingly, this analysis first exposed the heterogeneous integration landscapes between patients and the “preferred” chromatin state positions of integrants within patients (Fig. 6B). For example, Einkauf Patient 1 has a total of 42 integrants with the following positions: 2 (state 4, Strong transcription), 22 (state 5, Weak transcription), 1 (state 6, Genic enhancer), 8 (state 7, Active enhancer), 2 (state 8, ZNF genes & repeats), and 7 (state 15, Quiescent/low). Although most integrants are in state 5 (52.38%), state 7 (19.04%), and state 15 (16.66%), these encompass an extremely large portion of the genome (9.86%, 2.99%, and 73.84%, respectively) (Supplemental Table S5), thus these states are not significantly enriched for HIV proviruses.

For the three patients analyzed in the Einkauf dataset, two of them (Patients 1 and 3) showed nearly identical integration patterns, where HIV was preferentially enriched (35-fold and 51-fold enrichment for Patients 1 and 2, respectively) in state 8 (ZNF genes & repeats), which only represents a 0.14% fraction of the entire genome (Fig. 6B), and a minority of integrants in states 4-7 (for Patient 1) and states 2, 4, and 7 (for Patient 2). Despite this remarkable similar integration pattern in two patients, Patient 2 showed an enrichment of integrations in state 3 (Transcription TSS/TTS; 152-fold enrichment) and in state 6 (Genic enhancer; 19.8-fold enrichment), which only encompass 0.02% and 0.24% of the human genome, respectively (Supplemental Table S5).

A more highly diverse integration pattern was observed among the seven patients from the Kok dataset with Patient 4 almost exclusively containing positions in state 5 (Weak transcription), Patient 3 showing an enrichment of positions in state 7 (Active enhancer) and state 5 (Weak transcription), Patient 2 bearing an enrichment in state 6 (Genic enhancer), Patient 7 bearing an enrichment in state 4 (Strong transcription) and state 6 (Genic enhancer), and Patients 1, 5, and 6 containing HIV spread out in about 5-6 different states (Fig. 6B). Together, these studies illuminate the heterogeneous nature of HIV integration patterns in diverse chromatin states among patients, which has striking similarity to the immortalized (B-HIVE) system, possibly highlighting the plasticity in HIV transcriptional regulation leading to disparate proviral fates (silent-to-active).

### Persistence of Intact Versus Defective HIV Proviruses within Chromatin States

An important point is to be able to distinguish intact vs defective proviruses to start investigating where intact proviruses hide in the human genome and how they are regulated. To assess this, we interrogated the Einkauf dataset (Einkauf et al. 2019), which is the only study currently having this information for individual patients. Surprisingly, we observed a remarkable, almost completely distinct pattern of integration landscape enrichment between patients with intact proviruses in Patients 1 and 3 being enriched in state 8 (ZNF genes & repeats; 73-fold enrichment for Patient 1 and 76-fold enrichment for Patient 3), and intact proviruses in Patient 2 being enriched in state 3 (Transcription TSS/TTS; 208-fold enrichment) and a smaller enrichment in state 6 (Genic enhancer; 36-fold enrichment) (Fig. 6C and Supplemental Table S5), resembling the enriched HIV integration patterns in the immortalized system (B-HIVE) and other patient/*ex vivo* studies (Fig. 6A). Notably, a subset of integrants in the B-HIVE dataset (0.32%) contains examples that match the integration pattern of intact proviruses enriched in state 8 (ZNF genes & repeats) in the Einkauf dataset, again revealing conserved integration patterns in immortalized models of latency and primary/patient datasets.

In contrast to the enrichment of intact proviruses in Patients 1 and 3 in state 8 (ZNF genes & repeats), defective proviruses in these two patients were mainly enriched in state 4 (Strong transcription), state 5 (Weak transcription), and state 7 (Active enhancers), potentially indicating that intragenic and intra-enhancer positions, which typically correlate with high expression in the immortalized (B-HIVE) system, might be deleterious for replication competence. Given the rather small patient population and because the three patients have similar if not identical clinical and demographical characteristics (e.g., age, gender, CD4^+^ T cell counts, undetectable viral load, and time under suppressive therapy) (Einkauf et al. 2019), at this point it is impossible to predict what may be the cause of the variability in integration landscapes in these three patients.

Interestingly, an analysis of integration positions of intact vs defective proviruses in infants (GB) (Garcia-Broncano et al. 2019) revealed that intact proviruses were enriched in state 3 (Transcription TSS/TTS; 200-fold enrichment) while defective proviruses were enriched in state 7 (Active enhancer; 6.7-fold enrichment) and state 5 (Weak transcription; 2-fold enrichment) (Fig. 6C and Supplemental Table S5). Remarkably, the pattern of enrichment of intact proviruses in infants in the Garcia-Broncano dataset matches the enrichment of intact proviruses in adult Patient 2 of the Einkauf dataset potentially providing somewhat generalizable concepts of locations in the human genome in which intact proviruses persist. Similarly, the pattern of enrichment of defective proviruses in infants in the Garcia-Broncano dataset matches the enrichment of defective proviruses in adult Patients 1 and 3 of the Einkauf dataset potentially providing generalizable concepts of locations in the human genome in which defective proviruses persist.

## DISCUSSION

In this work, we have applied a previously unprecedented integrated approach: 1) to interrogate how the position of HIV proviruses in the human genome shape their expression, 2) to implement a machine learning approach to delineate features in the human genome predicting HIV proviral expression, and 3) to explore where defective and intact proviruses hide in the human genome and how their positions may dictate persistence in patients despite highly suppressive therapies, thereby regulating their future inducibility. Together, it is the combination of these three approaches that have enabled us to start defining relationships between HIV proviral positions in the human genome and their expression, to start understanding where intact HIV proviruses may be hiding to persist in patients, as well as to fuel questions for future studies.

Our studies on the exploration of HIV integration patterns and proviral expression revealed a number of salient points. First, B-HIVE insertions appear evenly on all chromosomes, with the majority of insertions occurring in genic regions (Fig. 2B), and that proviral expression is unrelated to the chromosome (Supplemental Fig. S1). Second, proviral expression has a slight correlation to distance to the most proximal human gene TSS when the insertion occurs inside a gene (Fig. 3A). Third, that both Intra- and Intergenic - Convergent orientation in relation to nearest gene has a slight correlation to proviral expression (Fig. 3B). Fourth, HIV expression has no sharp correlation with the distance to the most proximal TE; however, we observed a minor correlation between the position of HIV proviruses in Group 5, Intragenic - Convergent, and its distance to the most proximal SE (Fig. 3C). Fifth, mean HIV expression is higher in sub-compartment A1 compared to the other sub-compartments (A2, B1, B2, and B3; Supplemental Fig. S6G-H), which agrees with the higher enrichment of chromatin marks correlating with transcriptional activity and accessibility (Rao et al. 2014). Together, these studies suggest that HIV benefits from integration into more active compartments and with unique position and orientations respective to human genes and regulatory elements.

The ChromHMM analysis revealed that enhancers appear to dictate HIV proviral expression, potentially controlling its fate. For example, Active enhancers (state 7) and Bivalent enhancers (state 12) may provide a better environment for HIV expression compared to Quiescent/low genomic domains (state 15) (Fig. 4). Thus, while enhancers appear to be important regulators of HIV expression, their chromatin context, expression level at a given time and position in the 3D space may be more critical than the distance between the integration site and most proximal enhancer in the 1D space (Fig. 3C).

Our studies on the dissection of molecular features regulating HIV proviral expression prompted us to devise a machine learning model to predict features in the human genome contributing to proviral expression, which might be vital for intact proviruses persisting despite highly suppressive therapy. As such, the implementation of a logistic regression model trained with the B-HIVE integration/expression datasets showed good predictive power to be the first model employed to address this problem, with a sensitivity of 68.42% and a specificity of 59.10% (Fig. 5). This model revealed optimal features for the prediction, including upstream histone acetylation (H3K27ac) and chromatin accessibility (MNase-seq) and categorical nuclear sub-compartments. Strikingly, Cohn and colleagues described preferred integration in “hotspot” locations at or near repeat elements in patient samples (Cohn et al. 2015). This observation is interesting because our machine learning model only found H3K27ac and MNase-seq to be important, upstream relative to genome location of insert, which may be linked to the presence of active mobile elements in the human genome. Together, we envision that with the addition of larger datasets of proviruses, whose integration landscape and expression profiles are quantified with more robust technologies, could help to achieve newer models with better predictive powers. As the datasets grow, we expect that genetic landscape related to HIV proviral fate will be more prominent and, accordingly, future machine learning approaches can be used to provide a compass compatible with clinical decision-making.

Our studies on both the dissection and prediction of molecular features regulating HIV proviral expression and fate stimulated further data analyses to assess where HIV proviruses (both intact and defective) hide in the human genome and how their locations potentially regulate their expression. Since no single dataset yet exists to simultaneously interrogate HIV positions and expression in patient samples, we used a landmark dataset containing joint information on HIV positions and genome intactness (Einkauf et al. 2019). Given that intact proviruses can be expressed and thus reactivated to reseed the active viral reservoir upon cessation of therapy, we were mainly interested in comparing their positions in the human genome compared to defective proviruses. Interestingly, using this information and ChromHMM we showed that both intact and defective proviruses “hide” in different chromatin regions of the human genome (Fig. 6C), potentially suggesting that some areas are more permissive for “preserving” proviral intactness than others, which might be related to transcriptional and/or DNA replication effects. Strikingly, 2 out of 3 patients contained a significant fraction of enrichment of intact proviruses persisting in state 8 (ZNF genes & repeats; ∼4-fold) but also enriched about equally between states 1, 9,10 (Active TSS, Heterochromatin, and Quiescent/low, respectively; ∼3-5 fold). Furthermore, defective proviruses were preferably enriched, more than random chance, in states 4, 5, and 7 (Strong transcription, Weak transcription, and Active enhancer, respectively). Although intact proviruses were also detected in these chromatin states, as expected given their large coverage of the human genome, these positions do not differentiate proviral classes. Furthermore, an analysis of integration sites on infants also revealed that intact and defective proviruses were preferentially located in different chromatin states with intact proviruses primarily located in state 3 (Transcription TSS/TTS), and defective proviruses preferably located in states 7 (Active enhancers) and 5 (Weak transcription) (Fig. 6C). Remarkably, enrichment of defective proviruses in infants matches the enrichment of defective proviruses in adult Patients 1 and 3, but not Patient 2 (Fig. 6C).

While the patient data analysis revealed some interesting trends for the enrichment of intact vs defective proviruses in preferred locations (chromatin states), the lack of expression data does not allow us to interrogate relationships between their positions, expression, and inducibility. However, the combination of patient data analysis with the definition of features and chromatin states regulating HIV expression in the immortalized model of latency yielded a number of interesting observations that may be applicable to intact proviruses persisting in patient samples. For example, most of the intragenic proviruses, Groups 4-5 (convergent or same), are defective in patients, and intact HIV sequences in intergenic positions (divergent and convergent) are more prevalent than in genic positions, and/or close to TSS and/or to chromatin accessible areas (Einkauf et al. 2019). Also, while the transcription activity of both intragenic groups (Groups 4-5) is more correlated with the distance to the nearest TSS (Fig. 3A), these proviruses are typically defective in patients (Maldarelli et al. 2014; Einkauf et al. 2019), potentially due to their location inside genes and the high level of co-transcriptional processing.

Our studies spur some interesting ideas and questions for future research. Does the expression of HIV proviruses correlate with the activity of its neighborhood including any regulatory elements that can be transcribed such as promoters, enhancers, and repetitive elements? Does the directionality of HIV proviruses respective to most proximal genome regulatory elements (e.g., promoters, enhancers, repeats, etc.) matter? As such, we envision that future work will be needed: 1) to study differences in epigenomic landscapes before and after HIV insertion, 2) to collect larger integration libraries to study and develop better, more complete machine learning models, 3) to obtain more patient datasets with intactness for personalized medicine depending on where HIV inserts itself, and 4) to study HIV integrants at the single cell level to define their locations and intactness (MIP-seq), expression (scRNA-seq), and chromatin landscapes (scATAC-seq). Although these single-cell studies are the only ones that will allow to study the relationship between HIV position, intactness, and expression as well as viral-host chromatin landscapes, the scarcity of the latent reservoir in patients under suppressive therapy and the current inability to multiplexing these three technologies pose extreme limitations.

Given the above facts, exactly deciphering human genome codes shaping HIV proviral fate will certainly require painstaking investigations of multiple genetic and epigenetic features. To avoid incorrect assumptions that one or more regulatory features could contribute to position effects, deep and precise interrogations will have to be conducted in the same system, which will require the generation of thousands of clones containing single proviruses integrated into a unique position in the human genome and/or, even better, the use of single-cell level approaches to simultaneously characterize regulatory features of both HIV and human genomes, something that it is currently prohibited by the number of multi-omics approaches that can be implemented at the single cell level. Finally, our analysis demonstrates the importance of carefully and correctly characterizing enhancer elements when studying their potential role in genome regulation. Future studies should be cautious to employ the same rigorous standards when defining genomic domains to interrogate functional insights in human health and disease.

## METHODS

### NGS dataset analysis

Information on the library preparation and sequencing of MNase-seq as well as a description of data analysis for ChIP-seq, RNA-seq, TT-seq, MNase-seq, DNase-seq, and Hi-C can be found in the Supplemental Methods.

### Barcodes and HIV Integration Site Mapping on the Human Genome and HIV barcode clustering and quantification

B-HIVE data was processed with the B-HIVE for single provirus transcriptomics docker container (https://github.com/gui11aume/BHIVE_for_single_provirus_transcriptomics) with a change to the expr.nf file (see GitHub scripts for the updated script). The B-HIVE expression data set was subdivided into 6 different groups, 1) Intergenic – Same, 2) Intergenic – Convergent, 3) Intergenic – Divergent, 4) Intragenic – Same, 5) Intragenic – Convergent, and 6) Intragenic – Overlapping (which consists of 3 subgroups; Fig. 2A) depending on the relationship to the nearest gene from Gencode version 25. For each group, a Circos plot version 0.696 (Krzywinski et al. 2009) was created to show the relationship of HIV expression to genome location (Supplemental Fig. S1A-F). HIV expression versus distance to nearest TSS was plotted in R version R/3.3.2-gccmkl (Team 2014) using ggplot2 (Wickham 2016).

### Typical and Super enhancer Databases

Strand specific, TT-seq was used to identify possible typical and super enhancers. For this purpose, enhancers are defined as regions of the genome that are bi-directionally transcribed, and not in a gene nor its promoter, or an annotated linc-RNA. A 4 state HMM on TT-seq both strands, TT-seq forward strand, and TT-seq reverse strand identified regions of the genome that are actively transcribed. Four states were chosen over a 2-state model because there were various amounts of transcription found in the genome; genic regions were easily identified, but low expression intergenic regions could not be identified with 2 states. Thus, 3 of the 4 states were coded for transcribed, and 1 state was labeled as non-transcribed. A database of possible enhancers was created by identifying regions of the genome that were identified as transcribed for all data, and also overlapped with regions that were both forward and reverse transcribed. Also added to the possible enhancer list were regions where there were overlapping forward and reverse transcription, but not identified as transcribed in all data. From this list, protein-coding genes with 2 kb upstream and downstream were removed. Next, annotated regions from RNA-seq with an FPKM greater than 1 for all replicates and not protein-coding genes were removed. This final list of 20,943 regions is purported enhancers.

Super enhancers were identified using Rose v0.1 (Loven et al. 2013; Whyte et al. 2013), stitching together a 12.5 kb distance, excluding 2.5 kb from TSS. Purported enhancer regions from TT-seq were used as previously identified enhancer regions. Merged, filtered, and read mapped duplicates removed bam files of histone marks (H3K27ac, H3K4me3, and H3K4me1) were used to rank the possible enhancers. Since not all enhancers are identifiable with the 3-histone marks, TT-seq bam files were also used to identify super enhancers, and the output was filtered again for transcribed, annotated regions of the genome (e.g., genes and linc-RNAs). The final super enhancer database contained 767 merged regions, of which 360 were identified with H3K27ac alone, 436 identified with H3K4me3 alone, 301 identified with H3K4me1 alone, and 115 identified with TT-seq alone. The 767 super enhancer regions were removed from the 20,943 purported enhancer regions leaving 18,357 possible enhancer regions. Of these regions, 701 overlap with H3K27ac peaks, 262 overlap with H3K4me3 peaks, 702 overlap with H3K4me1 peaks, and 1301 overlap with forward and reverse TT-seq transcribed regions with bidirectional transcription. The final merged regions contain 2180 enhancers. The closest typical and super enhancers to each provirus within the 6 HIV insertion groups (Fig. 2A) were identified with bedtools closest (version 2.26.0) (Quinlan and Hall 2010). HIV expression versus distance to nearest typical enhancer or super enhancer was plotted in R version R/3.3.2-gccmkl (Team 2014) using ggplot2 (Wickham 2016).

### Identification of chromatin states – ChromHMM

We implemented ChromHMM (Ernst and Kellis 2017), which uses a multivariate (HMM) to calculate the probabilistic nature of a multi-state model and the biological nature of the state of chromatin at that location, in order to discover chromatin states in Jurkat T cells using epigenomics information derived from 7 individual ChIP-seq marks (H3K27me3, H3K4me3, H3K27ac, H3K4me1, H3K36me3, H3K79me3, and H3K9me3) known as state emissions. Filtered bam files, with mapping read duplicates removed, for each of the 7 histones, were individually converted to binary bin files with ChromHMM BinarizeBam v1.19 (Ernst and Kellis 2017). We first obtained the state emissions of 15 different chromatin states defined as described on the Roadmap epigenomics project (https://egg2.wustl.edu/roadmap/web_portal/chr_state_learning.html) on the basis of the observed data for the above 7 histone modifications. We used a core 15-state model for our analyses since it captured all the key interactions between the chromatin marks, and because larger numbers of states (e.g., “expanded 18-state model”) did not apparently capture sufficiently distinct interactions. To *de novo* generate the core 15-state model in Jurkat T cells we compared the relative abundance of the state emissions in Jurkat with known chromatin states for the 3 ENCODE cell lines most genetically and phenotypically linked with Jurkat (E115: Dnd41 T cell leukemia, E116: GM1282878 lymphoblastoid, and E123: K562 T cell leukemia) (Supplemental Fig. S3A-C). In order to assign biologically meaningful mnemonics to the 15 chromatin states, we used the ChromHMM package to compute the overlap and neighborhood enrichments of each chromatin state relative to various types of functional annotations including the ChromHMM built in RefSeq annotations of: 1) CpG islands, 2) genes, 3) exons, 4) introns, 5) TSS, 2-kb windows around TSS (TSS flanking), transcription termination sites (TTS), and 2-kb windows around TTS (TTS flanking) based on the GENCODE v27 annotation, 6) Zinc finger (ZNF) genes obtained from ChromHMM, and 7) typical enhancers and super enhancers obtained as described above (Fig. 4A).

### Identification of sub-compartments

The two Jurkat Hi-C libraries (see Supplemental Methods) were combined with HOMER v4.10.4 makeTagDirectory using a standard protocol (Heinz et al. 2010). The eigen values of the first principle component of each chromosome was calculated with HOMER v4.10.4 runHiCpca using a standard protocol (Heinz et al. 2010). The sign of the eigen values divide each chromosome; however, it does not state if either positive or negative values are representative of the A or B sub-compartments. So, for each chromosome, the positive and negative eigen values were overlaid with Jurkat’s 15-state chromatin marks (ChromHMM v1.19 OverlapEnrichment) (Ernst and Kellis 2017). The A and B sub-compartments were clearly defined with the B sub-compartments preferentially segregating in states 9, 13, and 15 (heterochromatin, repressed Polycomb, and quiescent/Low, respectively). The sign of the eigen values were then corrected so that positive values represented the A sub-compartment and negative values, B sub-compartment. K-means of k = 2 was calculated on the A sub-compartment, and k-means k = 2 through k = 5 on the B sub-compartment using R v3.5.1 k-means. The A sub-compartment was labeled A1 or A2 based on the results of overlaying the k-means on the chromatin states; with A1 having much higher values than A2. The B sub-compartment was overlaid with the chromatin states for all k-mean k = 2 through k = 5, however, the results did not split the B sub-compartment on expected chromatin marks (Supplemental Fig. S5). Thus, the B sub-compartment was not sub-divided. The GM12878 sub-compartments were retrieved from the GEO database (GSE63525). The coordinates were lifted over to GRCh38 with UCSC liftOver (Hinrichs et al. 2006).

### Machine learning

In order to study the immediate landscape surrounding HIV insertions (1559 insertions in total) and its possible effects on expression, we looked at 2-kb regions, in 200-bp increments around the HIV integration sites (Fig. 5A). RPKM values of each 200-bp bin (20 bins in total) were calculated for the 7 histone marks (H3K27ac, H3K4me3, H3K4me1, H3K36me3, H3K79me3, H3K9me3, and H3K27me3), RNA-seq, MNase-seq, DNase-seq, and TT-seq using RPKM.py (https://git.biohpc.swmed.edu/venkat.malladi/miscellaneous_scripts/blob/master/scripts/rpkm.py). Discrete values for ChromHMM states and lamin sub-compartment states were also noted for each 200 bp region. The ChromHMM states were then converted from a categorical into a numerical value based on our understanding on its openness: U1 (most open), U4, U3, U6, U2, U7, U5, U10, U8, U11, U12, U14, U13, U9, and U15 (most close) in order. The HIV expression level was normalized by z-transform and was annotated as ‘Low’ if the normalized expression is lower than −0.5 (n=351), as ‘High’ if higher than 0.5 (n=455), and otherwise as ‘Intermediate’ (n=753) (Fig. 5A). To determine optimal features, which have predictive power in HIV fate prediction, and train a prediction model with them, a Machine Learning (ML) approach was taken. As a first step, the genetic landscape dataset was randomly split into a training dataset (75% of HIV insertions) and a test dataset (25% of HIV insertions). To select optimal features for HIV expression level prediction, an R package, *smbinning* (https://rdrr.io/cran/smbinning/), was applied to the training dataset consisting of ‘High’ and ‘Low’ expression instances only. It returned each feature’s Information Value (IV), which is relevant to its importance in the prediction task, and features of IV ≥ 2 were determined as optimal ones (Supplemental Table S2). After training a logistic regression model with the training dataset of the optimal features, the trained model was evaluated with the unseen test dataset. Note that ‘Intermediate’ expression instances were excluded for the model training to get a better model, but included for the model evaluation. A metagene profile plot of 2 kb region surrounding HIV insertion in H3K27ac and MNase-seq (Supplemental Fig. S7A,B) as well as heatmaps of HIV insertions and mean expression in each sub-compartment (Supplemental Fig. S7C,D) were created.

### Patient Data

Patient data was downloaded (Supplemental Table S4) and, if necessary, lifted over from older assemblies to GRCh38 using UCSC liftOver (Hinrichs et al. 2006). Heatmaps were created with ChromHMM OverlapEnrichment (Ernst and Kellis 2017).

## DATA ACCESS

### URLs

The computer code used for this analysis is available on GitHub: https://github.com/utsw-bicf/HIVproviral_fate

### Data availability

NGS datasets used in this study (Supplemental Table S1) were downloaded from NCBI Gene Expression Omnibus (www.ncbi.nlm.nih.gov/geo/; (Barrett et al. 2013) or ENCODE project (www.encodeproject.org).

### Data deposition

MNase-seq, raw and processed sequencing results, have been submitted to NCBI Gene Expression Omnibus (GEO; https://www.ncbi.nlm.nih.gov/geo/) under accession number GSE144753.

## ACKNOWLEDGEMENTS

We thank members of the D’Orso laboratory (Nora-Guadalupe Ramirez, Ashutosh Shukla, Jinli Wang, and Usman Hyder) for critical reading of the manuscript and Chi Pak for generating the MNase-seq dataset. We are indebted to Mathias Lichterfield for sharing the patient datasets from the Einkauf and Garcia-Broncano studies as well as the many groups whose datasets were interrogated in this study. This research was supported in part by US National Institutes of Health under award number R01AI114362 to I.D. and by the Cancer Prevention Research Institute (CPRIT) under award number RP150596 to H.R. and J.L. The content is solely the responsibility of the authors and does not necessarily represent the official views of the US National Institutes of Health.

## AUTHOR CONTRIBUTIONS

H.R. and I.D. developed the idea. H.R., J.L., and I.D. designed the experiments. H.R., J.L., and I.D. analyzed the data. H.R. and I.D. wrote the manuscript with feedback from J.L., C.G. and V.M.

