## Supplemental Figure and Methods for "An integrated genomics approach towards deciphering human genome codes shaping HIV-1 proviral transcription and fate"

### SUPPLEMENTAL FIGURES

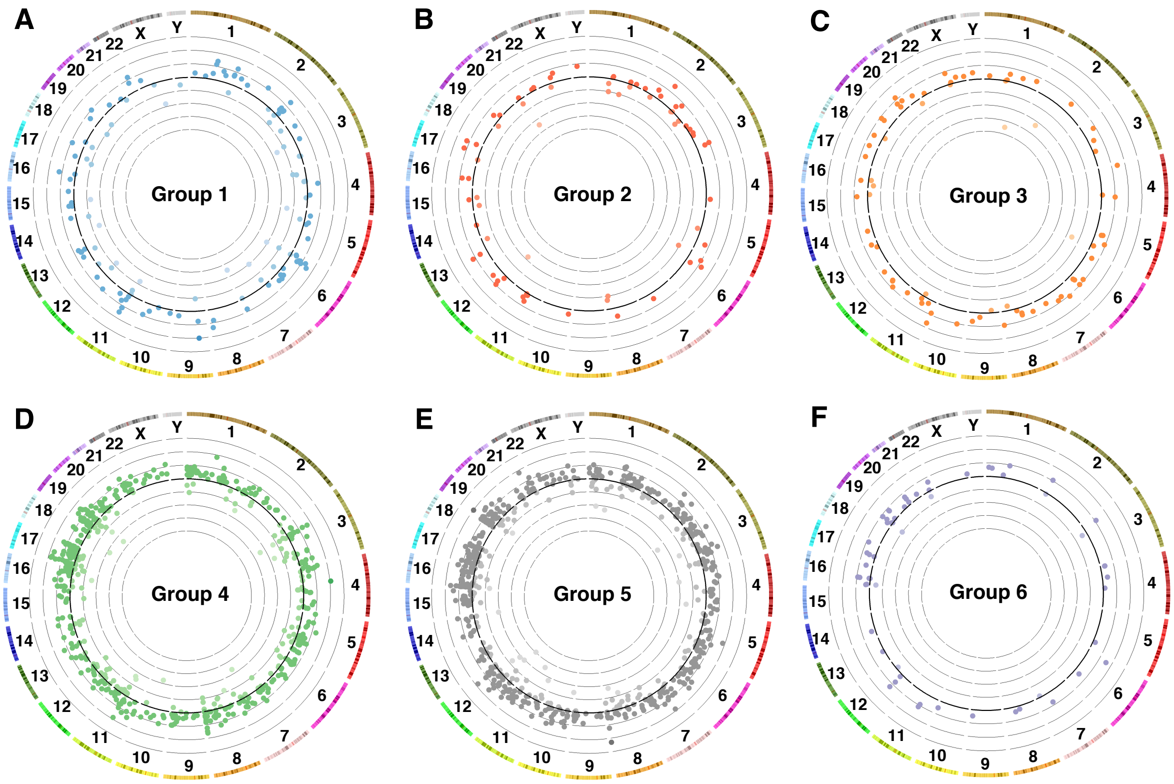


**Supplemental Figure S1.** Circos plots of proviral integration and expression for HIV integration groups classified based on their positions and orientations respective to human genes. (A-F) Circos plot of each of the 6 HIV integration groups as described in Fig. 2A. Group1: Intergenic - Same, Group 2: Intergenic - Convergent, Group 3: Intergenic - Divergent, Group 4: Intragenic - Same, Group 5: Intragenic - Convergent, and Group 6: Intragenic - 2 Genes Overlapping. Each circle represents B-HIVE chromosomal distribution and expression, with the inner most line, a $\log_{10}$ of HIV expression = -4, and the outer most line, a $\log_{10}$ of HIV expression = 3.

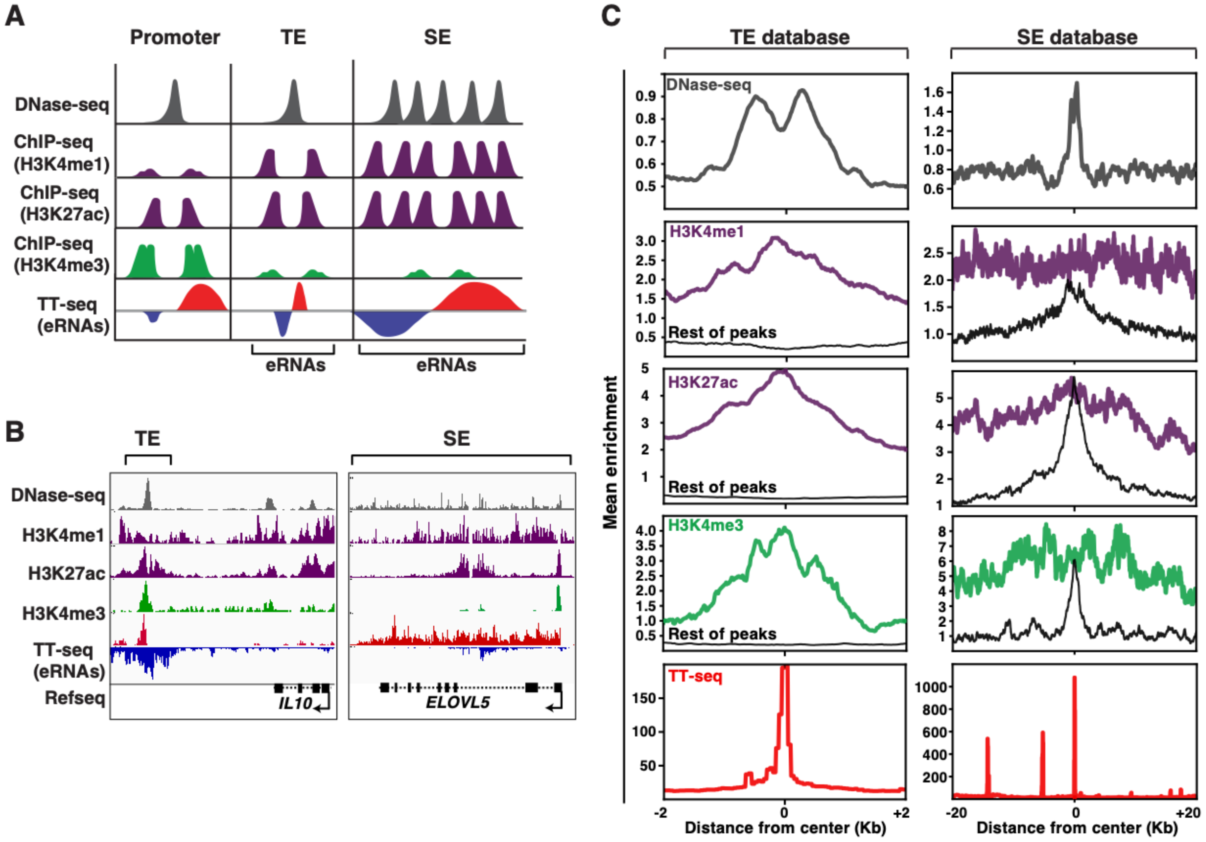


**Supplemental Figure S2.** Identification of enhancers with nascent transcripts and histone marks. (A) Features of promoters and enhancers [typical (TE) and super (SE)]. Promoters and enhancers are characterized by the presence of DNase I hypersensitive sites (DNase-seq) and H3K27ac-modified chromatin. However, compared to promoters, enhancers contain greater levels of H3K4me1-modified chromatin, and lower levels of H3K4me3-modified chromatin. Promoters, TE, and SE support bidirectional transcriptional activity, but enhancers produce a class of bidirectional and symmetric non-coding RNAs (eRNAs), whereas promoters support asymmetric transcriptional activity (i.e., higher transcription levels are observed in the coding strand). (B) Genome browser track view of a representative TE and SE in Jurkat T cells. (C) Metagene plots of DNase-seq, various epigenetic markers, and nascent transcriptional activity (TT-seq) for TE and SE in Jurkat T cells. For histone marks, TE and SE are further subdivided into enhancers that were identified with those histone marks and enhancers identified by all other marks (rest of peaks).

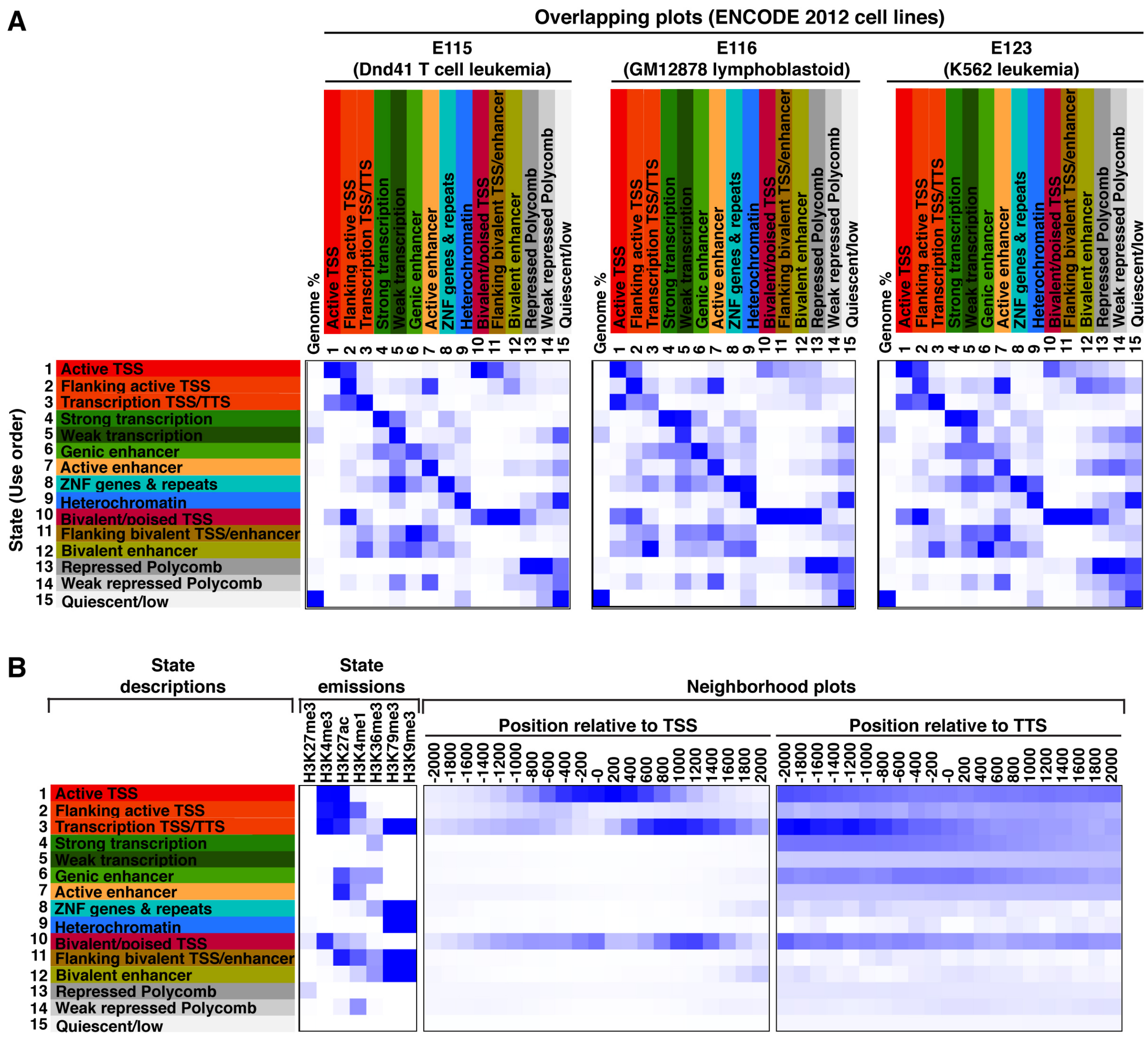


**Supplemental Figure S3.** ChromHMM heatmaps. (A) Overlap enrichment plots (ChromHMM) comparing Dnd41 T-cell leukemia (E115), GM12878 lymphoblastoid (E116), and K562 leukemia (E123) 15 state marks with Jurkat T cell 15 state marks. (B) Neighborhood and emissions plots of the Jurkat T cell 15 state marks.

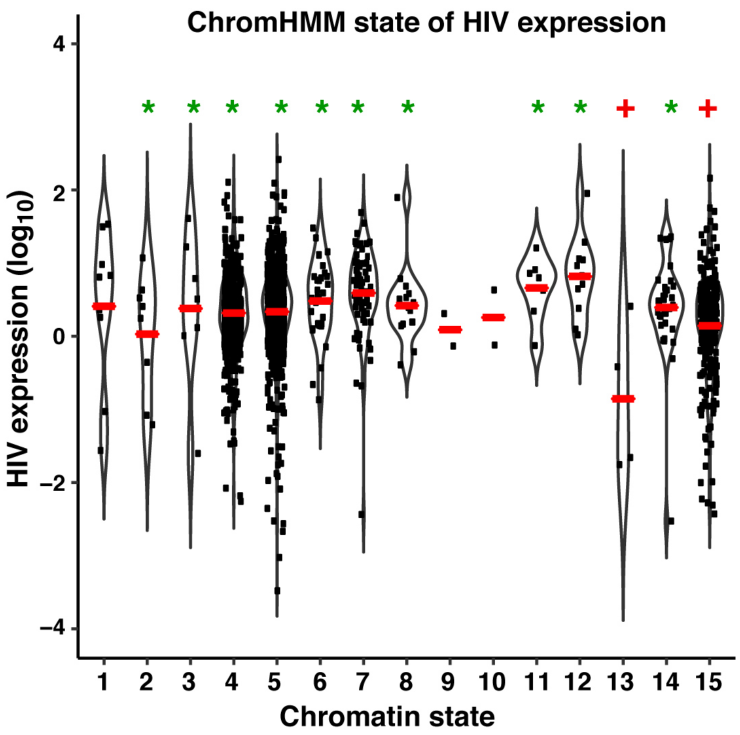


**Supplemental Figure S4.** B-HIVE insertions and expression by Jurkat T cell 15-state model. Jitter and violin plot of B-HIVE insertions for each of the Jurkat T cell 15 chromatin states. Green asterisks are states where insertions are more than expected by random occurrence ($P$ < 0.05, two-proportions z-test). Red crosses are states where there are significantly less insertions than expected ($P$ < 0.05, two-proportions z-test). The red line represents mean $\log_{10}$ HIV expression for each chromatin state.

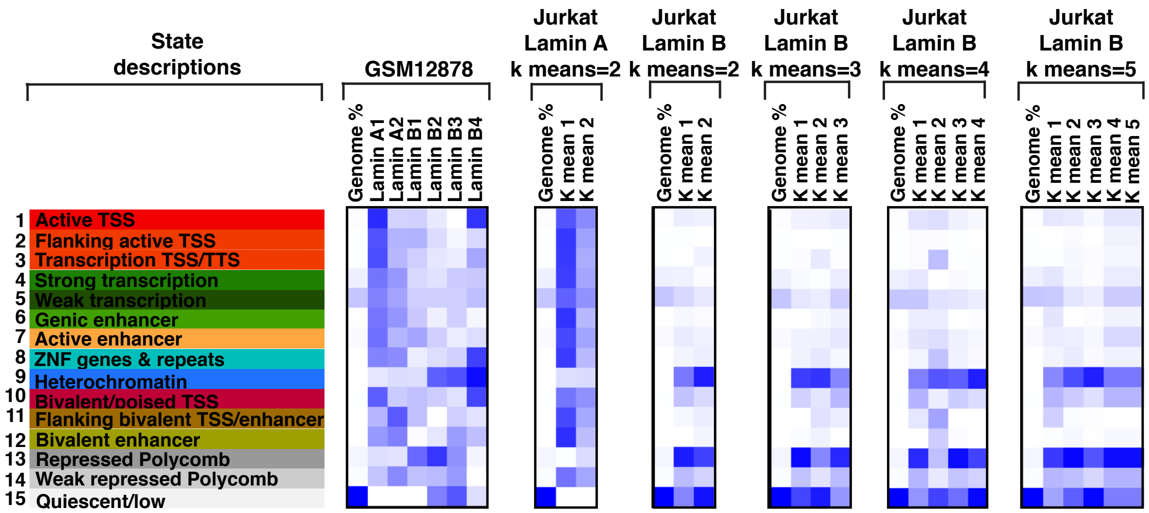


**Supplemental Figure S5.** K-means clustering of Jurkat T cell Hi-C data. From left to right: 15-state Jurkat T cell model. Overlap enrichment plot (ChromHMM) of GSM12878 in Jurkat T cell 15 states. K-means clustering of 2 for the A sub-compartment from the first eigenvector of Jurkat T cell Hi-C data. K-means clustering of 2 through 5 for the B sub-compartment from the first eigenvector of Jurkat T cell Hi-C data.

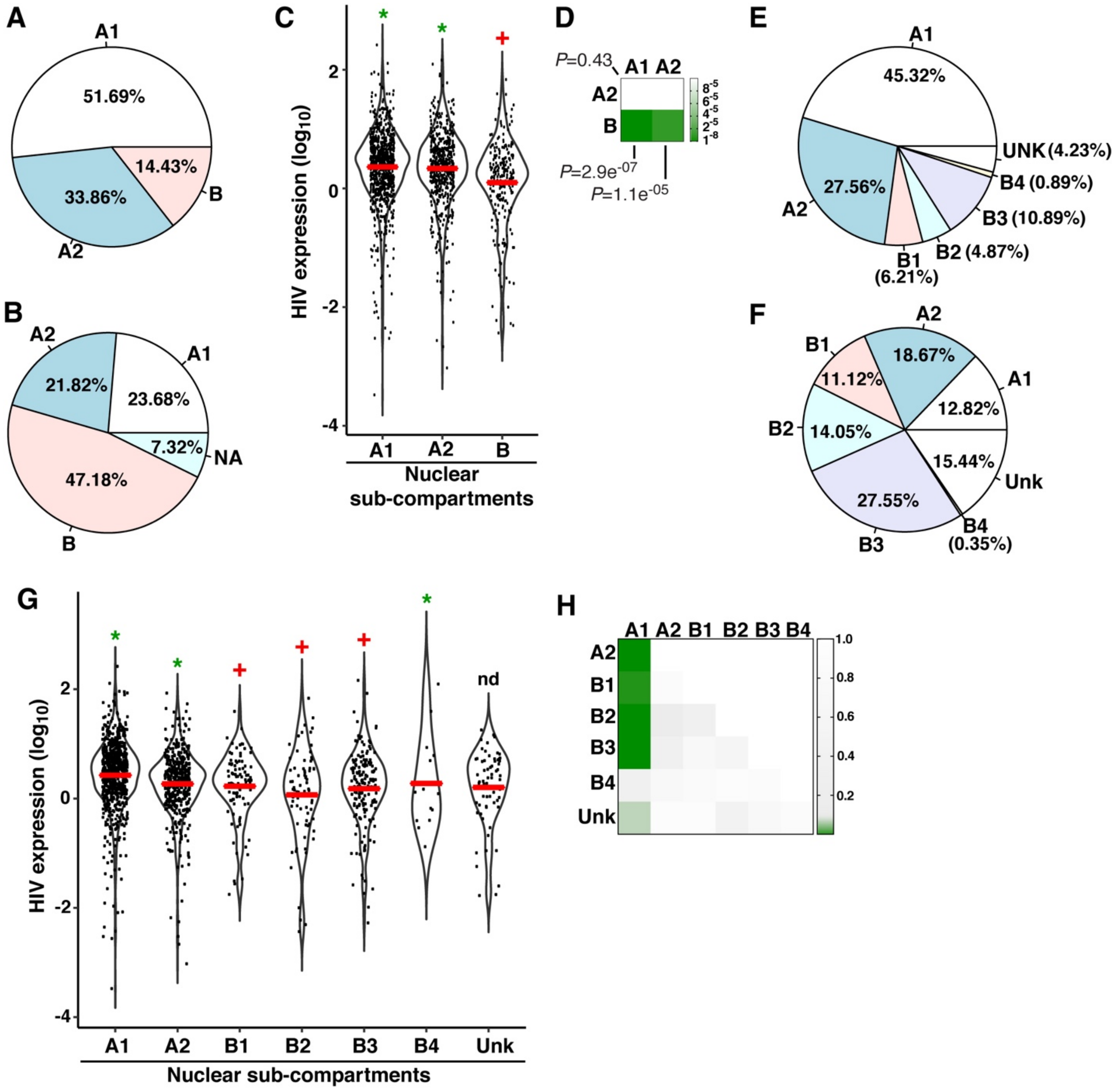


**Supplemental Figure S6.** Expression of HIV integration and expression in relation to 3D architecture. (A) Pie chart of HIV integration per Jurkat T cell sub-compartments derived from Hi-C data. (B) Pie chart of Jurkat T cell sub-compartment genomic coverage derived from Hi-C data. (C) Jitter and violin plot of HIV expression in each Jurkat T cell sub-compartments derived from Hi-C data. The red line represents mean expression. Green asterisks represent significantly more insertions relative to sub-compartment genomic coverage; red crosses represent significantly less insertions. (D) Heatmap representing the p-value (*P*) pairwise comparison of sub-compartments derived from Jurkat T cell Hi-C data vs expression using a Kruskal-Wallis rank sum test. (E) Pie chart of HIV integration per GM12878 nuclear sub-compartments. Unk denotes unknown sub-compartments. (F) Pie chart of GM12878 sub-compartments genomic coverage. Unk denotes unknown sub-compartments. (G) Jitter and violin plot of HIV expression in each GM12878 sub-compartments derived from Hi-C data. The red line represents mean expression. Green asterisks represent significantly more insertions relative to sub-compartment genomic coverage, and red crosses represent significantly less insertions; nd was not calculated due to unknown (Unk) sub-compartments. (H) Heatmap representing the p-value (*P*) pairwise comparison of sub-compartments derived from GM12878 vs expression using a Kruskal-Wallis rank sum test.

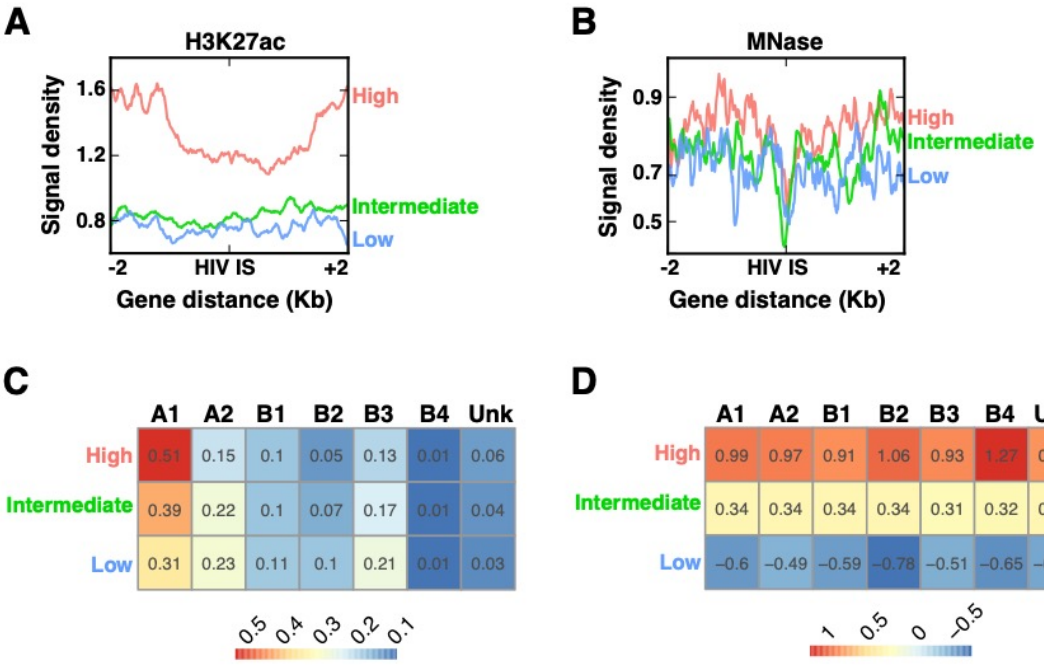


**Supplemental Figure S7.** Plots of optimal features surrounding HIV insertion versus expression level prediction based on surrounding genetic landscape. (A-B) Metagene plots of optimal features (H3K27ac and MNase-seq) 2-kb surrounding B-HIVE insertions, split by the three categories based on expression (Low, Intermediate, and High). (C) Heatmap of the percentage of B-HIVE insertions, split by the three categories of Low, Intermediate, and High, versus GSM12878 sub-compartments. (D) Heatmap of the mean expression of B-HIVE insertions, split by the three categories (Low, Intermediate, and High) versus GSM12878 sub-compartments.

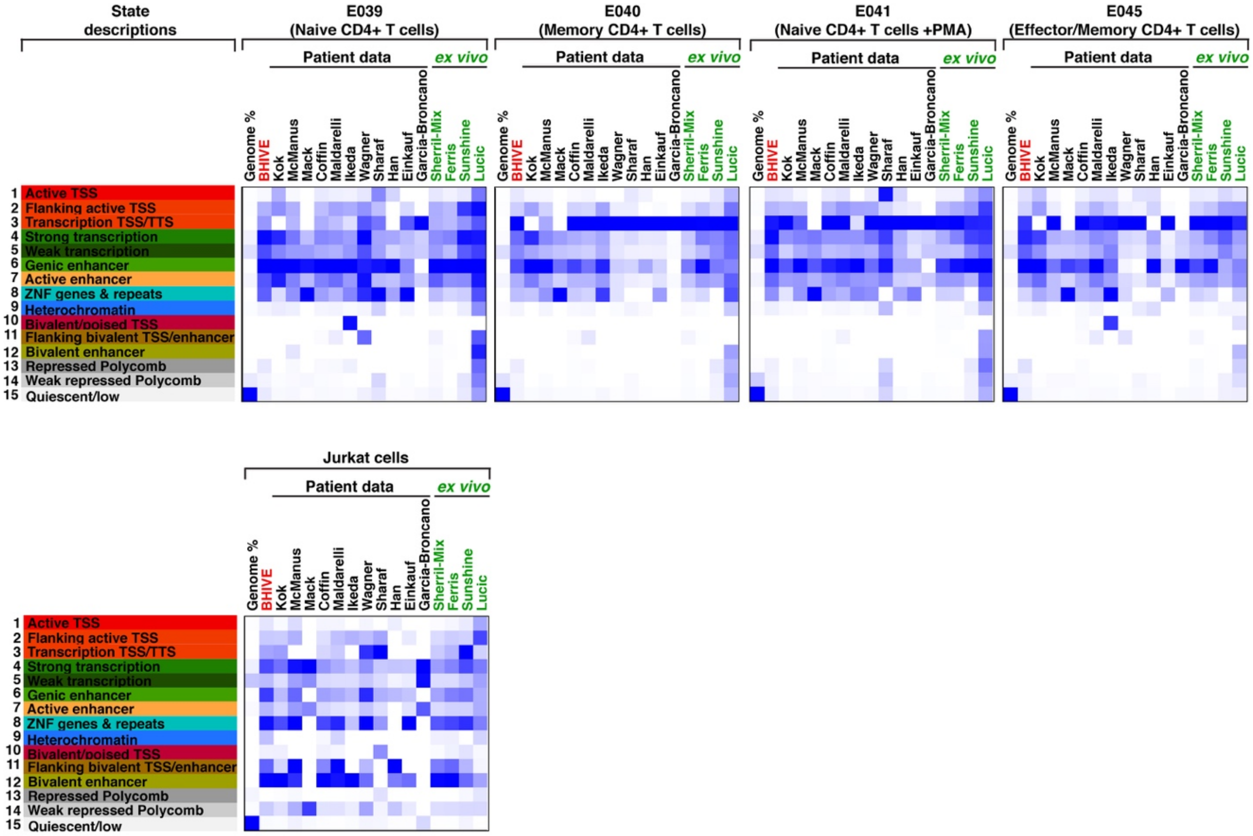


**Supplemental Figure S8.** Heatmaps of patient and *ex vivo* data on 4 primary CD4$^{+}$ T cell states and Jurkat T cells. Overlap enrichment plots (ChromHMM) of 11 studies involving patient and 4 studies in *ex vivo* compared to each of the indicated primary CD4$^{+}$ T cell states (top) and Jurkat T cells (bottom).

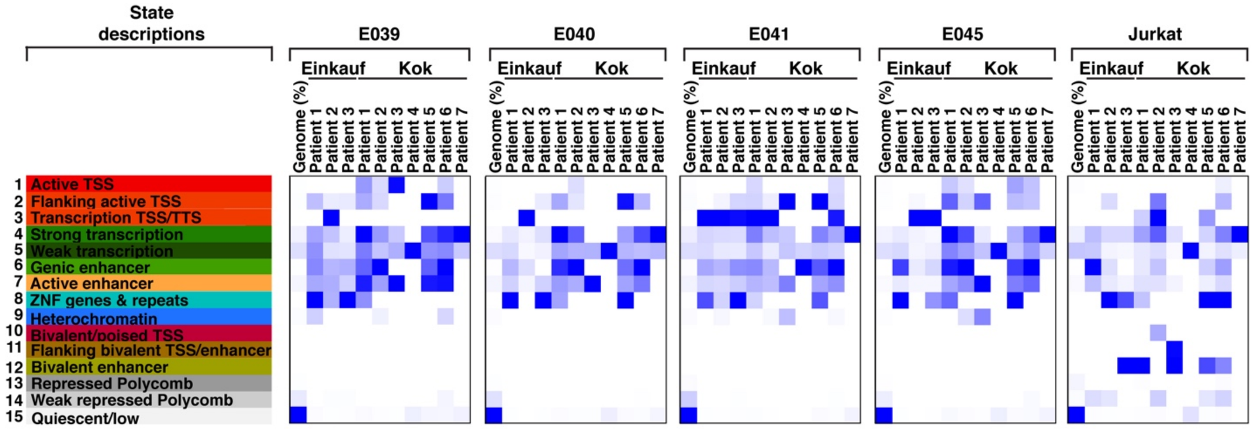


**Supplemental Figure S9.** Heatmaps of individual patient data on 4 primary $CD4^+$ T cell states and Jurkat T cells. Overlap enrichment plots (ChromHMM) of Einkauf and Kok, split by individual patients.

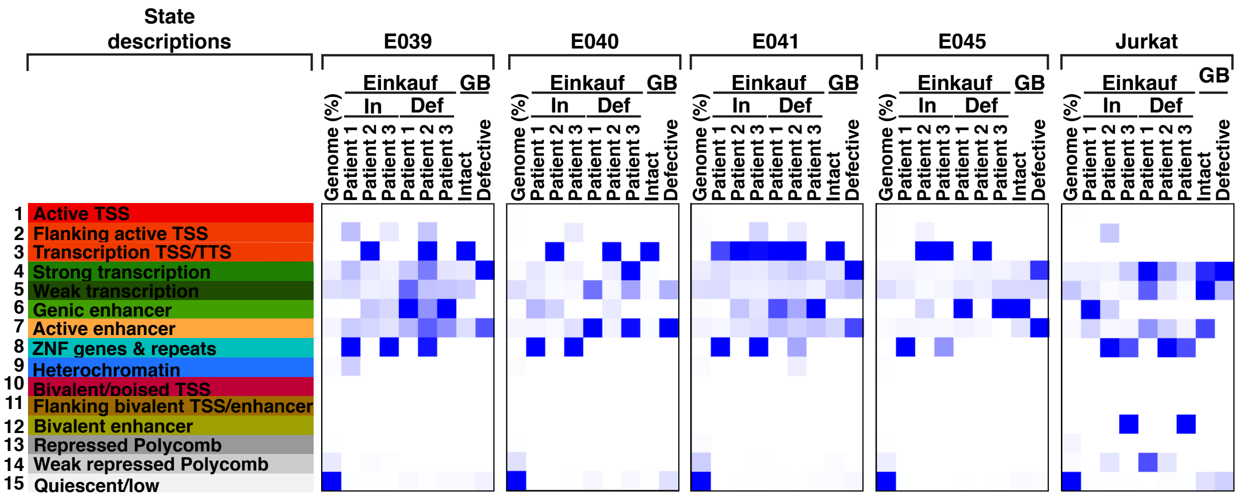


**Supplemental Figure S10.** Heatmaps of intact or defective proviruses of patient data on 4 primary CD4$^{+}$ T cell states and Jurkat T cells. Overlap enrichment plots (ChromHMM) of integration positions of patients from the Einkauf and Garcia-Broncano (GB) datasets, split by intact and defective proviruses.

### SUPPLEMENTAL METHODS

#### ChIP-seq data analysis.

The Nextflow (Di Tommaso et al. 2017) BICF ChIP-seq Analysis Workflow version 1.0.0 (Barnes et al. 2019) processed all ChIP-seq files, merging separate experiments as technical replicates. Briefly, reads were trimmed with trimgalore version 0.4.1 (Martin 2011) (parameters: -q 25 --illumina --gzip --length 35), aligned with bwa aln (-q 5 -l 32 -k 2) and then bwa samse (standard parameters) version 0.7.12 (Li and Durbin 2009), sorted and indexed with SAMtools version 1.3 (Li et al. 2009) (-F 1804 -q 30), and duplicates removed with Sambamba version 0.6.6 (Tarasov et al. 2015) (standard options). Bam files were converted to tagAlign with bedtools version 2.26.0 bamtobed (Quinlan and Hall 2010), after which, samples were checked for quality control using deeptools version 2.5.0.1 multiBamSummary, plotCorrelation, plotCoverage, and plotFingerprint (all standard protocols) (Ramirez et al. 2016) and cross correlation analysis with phantompeakqualtools version 1.2 (Kharchenko et al. 2008; Landt et al. 2012). Peaks were called with MACS2 version 2.1.0-20151222 (Zhang et al. 2008), using the predominant fragment length from the cross correlation analysis as --extsize (other parameters: -p 1e-2 --nomodel --shift 0 --keep-dup all -B --SPMR). Consensus peaks were called (bedtools version 2.26.0; (Quinlan and Hall 2010) and annotated (library ChIPseeker in R; Team 2014; Yu et al. 2015) if at least 2 replicates or pseudo-replicates contained a peak.

#### RNA-seq data analysis.

FASTQ files were processed with the BICF RNA-seq Analysis workflow version 0.5.5. Briefly, reads with phred quality scores less than 20 and less than 35 bp after trimming were removed from further analysis using trimgalore version 0.4.1 (Martin 2011). Quality-filtered reads were then aligned to the human reference genome (GRCh38) using the HISAT version 2.0.1 (Pertea et al. 2016) aligner using default settings and marked duplicates using Sambamba version 0.6.6 (Tarasov et al. 2015). Aligned reads were quantified to coding sequences of known transcripts using ‘featurecount’ version 1.4.6 (Liao et al. 2014) per gene ID against Gencode version 25. HIV expression versus $\log_{10}$ Fragments Per Kilobase Million (FPKM) of the nearest gene (e.g., the gene in which HIV is integrated into if intragenic or the nearest gene if intergenic) was plotted in R version R/3.3.2-gccmkl (Team 2014) using ggplot2 (Wickham 2016).

#### TT-seq data analysis.

FASTQ files were processed a modified version of the BICF RNA-seq Analysis workflow version 0.5.5. Briefly, reads with phred quality scores less than 20 and less than 35 bp after trimming were removed from further analysis using trimgalore version 0.4.1 (Martin 2011). Quality-filtered reads were then aligned to the human reference genome (GRCh38) using the HISAT version 2.0.1 (Pertea et al. 2016) aligner using default settings and marked duplicates using Sambamba version 0.6.6 (Tarasov et al. 2015). Aligned reads were quantified to the entire annotated transcript region using ‘featurecount’ version 1.4.6 (Liao et al. 2014) per gene ID against Gencode version 25.

#### MNase-seq library preparation.

Jurkat $CD4^+$ T cells were cultured in RPMI 1640 media supplemented with 10% Fetal bovine Serum (FBS) and 1X Penicillin/Streptomycin at 37°C with 5% $CO_2$ at optimal density of 0.5-1 x $10^6$ cells per mL. Cells were passaged every 2 days at a 1/3 dilution. Cell suspensions were transferred to 50 mL conical tubes and pelleted at 420 x *g* for 10 min. Cells were then resuspended in PBS at a density of 1 x $10^6$ cells/mL for crosslinking with 0.5% methanol-free formaldehyde (ThermoFisher) at room temperature with rotation for 10 min. The reaction was then quenched with 150 mM glycine (PBS buffer pH 7.5) for 10 min at room temperature. Cells were then pelleted by centrifuging at 420 x *g* for 10 min at 4°C and then washed twice with 20 mL cold PBS each time. Nuclei were collected by lysing the cells in Farnham's lysis buffer (5 mM PIPES pH 8.0, 85 mM KCl, and 0.5% NP-40 freshly supplemented with 1 mM PMSF and EDTA-free Protease inhibitor cocktail), washed once with cold MNase buffer (20 mM Tris-HCl pH 7.5, 15 mM NaCl, 60 mM KCl, and 2 mM $CaCl_2$) and then resuspended in MNase buffer at a concentration of 10 x $10^6$ cells/mL before digestion. Micrococcal nuclease (New England BioLabs, M0247S) digestion was performed using 1100 U enzyme/$10^6$ cells at 37°C for 10 min to achieve roughly 80% mono-nucleosome – 20% di-nucleosome populations. The reaction was interrupted with Stop buffer (20 mM EDTA pH 8.0, 20 mM EGTA pH 8.0, and 0.4% SDS) and then centrifuged at 21,000 x *g* for 5 min at 4°C. The supernatants were saved, and the small white pellet discarded. Samples (100 µL) were mixed with 1 volume of 2X Proteinase K buffer (4 mM EDTA pH 8.0, 40 mM Tris-HCl pH 6.8, 1M NaCl, and 1 mg/mL Proteinase K) for reverse crosslinking at 65°C for 16 hrs. After reverse crosslinking the samples were first extracted with 1 volume of phenol-chloroform-isoamyl alcohol (25:24:1 ratio) with centrifugation at 21,000 x *g* for 5 min at 4°C and later with 1 volume of chloroform-isoamyl alcohol (24:1 ratio). The aqueous phase was transferred to a 1.5 mL epitube and precipitated with 2.5 volumes of 100% cold Ethanol and 1/10 volume of 3M NaOAc with centrifugation at 21,000 x g for 15 min at 4°C. Samples were finally washed with 75% Ethanol, air dried for ~5 min and resuspended in 20 µL of water. DNA concentration was measured by Qubit/Nanodrop. About 3 µg of DNA were loaded onto a 1.5% DNA agarose gel to verify the expected nucleosomal size distribution (80% mono- and 20% di-nucleosome). The mono-nucleosome band (~150 bp size) was excised and gel cleaned up using DNA Clean & Concentrator kit (Zymo Research) following the manufacturer's instructions. DNA was eluted with 20 µL water (25 ng/µL final concentration). Replicate DNA samples were analyzed on high sensitivity DNA tape on Agilent 2200 TapeStation and used for library preparation. Library was prepared with ~375 ng of mono-nucleosomal DNA using the KAPA Hyper Prep Kit (KAPA Biosystems, KK8502) according to the manufacturer's instructions. For the PCR amplification step, we inputted ~7.5 ng and performed 9 cycles of amplification, obtaining 700 ng (~35 ng/µL). The quality control of the MNase-seq library was done on Agilent 2200 TapeStation. A single peak with average size of 306 bp (including ligated adapters) was observed. The MNase-seq DNA library was diluted to 4.2 nM for sequencing on an Illumina NextSeq 500 instrument as a 2 x 75 bp library. Illumina bcl2fastq (v 2.19.0) software was used for basecalling.

**MNase-seq data analysis.**
FASTQ files were processed with a modified Nextflow (Di Tommaso et al. 2017), BICF ChIP-seq Analysis Workflow version 1.0.0 (Barnes et al. 2019). Briefly, we used trimgalore version 0.4.1 (Martin 2011) on the raw reads to remove reads shorter than 35 bp and with phred quality scores less than 20 and then aligned trimmed reads to the human reference genome (GRCh38) using default parameters in BWA sample version 0.7.12 (Li and Durbin 2009). The aligned reads were subsequently filtered for quality and uniquely mappable reads were retained for further analysis using SAMtools version 1.3 (Li et al. 2009) and Sambamba version 0.6.6 (Tarasov et al. 2015), and bedtools version 2.26.0 (Quinlan and Hall 2010) bamtobed converted the bed file to tagalign. Peaks were called with iNPS version 1.2.2 (Chen et al. 2014) and filtered for a $-\log_{10}$ (Pvalue_of_peak) of less than 0.05.

**DNAse-seq data analysis.**
FASTQ files were processed with a modified Nextflow (Di Tommaso et al. 2017), BICF ChIP-seq Analysis Workflow version 1.0.0 (Barnes et al. 2019). Briefly, we used trimgalore version 0.4.1 (Martin 2011) on the raw reads to remove reads shorter than 35bp and with phred quality scores less than 20 and then aligned trimmed reads to the human reference genome (GRCh38) using default parameters in BWA samse version 0.7.12 (Li and Durbin 2009). The aligned reads were subsequently filtered for quality and uniquely, mappable reads were retained for further analysis using SAMtools version 1.3 (Li et al. 2009) and Sambamba version 0.6.6 (Tarasov et al. 2015).

Relaxed peaks were called using MACS2 version 2.1.0-20151222 (Zhang et al. 2008) with the following parameters: -p 1e-2 --nomodel --shift -100 --extsize 200 --keep-dup all -B --SPMR. Peaks that overlap at least 50% between replicates were retained.

**Hi-C data analysis.**

Reads were pooled by library and ran through the standard Hi-C pipeline using HOMER version 4.10.4 (Heinz et al. 2010). Briefly, reads were trimmed with homerTools trim -3 GATC -mis 0 -matchStart 20 -min 20, mapped to human reference genome (GRCh38, canonical) with bowtie2 version 2.2.8 (Langmead and Salzberg 2012), and converted to tag directory (makeTagDirectory -genome hg38 -checkGC -restrictonSite GATC). Matrices are normalized with analyzeHiC (standard protocol).
